# E3 Ubiquitin Ligase Highwire/Phr1 Mediates Endocytic Control of JNK Signaling in *Drosophila* Neurons

**DOI:** 10.1101/2024.11.03.621782

**Authors:** Srikanth Pippadpally, Anjali Bisht, Saumitra Dey Choudhury, Manish Kumar Dwivedi, Zeeshan Mushtaq, Leishangthem Biteshwari Devi, Suneel Reddy-Alla, Vimlesh Kumar

**Affiliations:** Department of Biological Sciences, Indian Institute of Science Education and Research (IISER) Bhopal, Academic Building 3, Bhauri, Bhopal-462 066, Madhya Pradesh, India; College of Community Science, Acharya N.G. Ranga Agricultural University, Guntur-522034, Andhra Pradesh, India; Centralized Core Research Facility (Microscopy) Convergence Block, All India Institute of Medical Sciences (AIIMS), Ansari Nagar, New Delhi-110029, India

**Keywords:** Endocytosis, BMP pathway, JNK pathway, Highwire assemblies, Wallenda, Rab11, Synaptic growth signaling

## Abstract

Endocytosis is essential for synaptic membrane retrieval and the trafficking of growth-regulatory signaling components, yet the mechanistic links between endocytic dysfunction and synaptic overgrowth have remained elusive. Here, we uncover a previously unrecognized mechanism by which endocytosis constrains synaptic growth by regulating the neuronal E3 ubiquitin ligase Highwire (Hiw/Phr1), a key negative regulator of the c-Jun N-terminal Kinase (JNK) signaling pathway. We show that loss of endocytic proteins causes Hiw/Phr1 to accumulate in the neuronal cell bodies, leading to elevated MAP3K Wallenda (Wnd)/DLK levels and hyperactivation of JNK signaling. The accumulated Hiw/Phr1 assembles into dynamic punctate structures, as revealed by their rapid and reversible dissolution with 1,6-hexanediol (1,6-HD). Acute blockade of endocytosis using a conditional temperature-sensitive dynamin mutant *Shibire^ts^*similarly triggered robust Hiw accumulation at restrictive temperatures. We further demonstrate that Rab11-positive recycling endosomes are essential for proper Hiw localization and turnover, directly linking endosomal trafficking to the control of JNK signaling. Finally, we show that both the Bone Morphogenetic Protein (BMP) pathway and JNK signaling are necessary and sufficient to guide synaptic morphogenesis at the *Drosophila* neuromuscular junction (NMJ), thereby integrating endocytic trafficking with synaptic growth signaling. Our findings establish endocytosis as a critical regulator of Hiw/Phr1-dependent JNK signaling, with implications that extend beyond synaptic morphogenesis to axon injury and degeneration pathways.

## Introduction

Synapse formation, elimination, and remodeling are fundamental processes in neuronal circuit development and are essential for dynamic plasticity that enables adaptation to sensory experience and physiological stress (BASTRIKOVA *et al*. 2008; COLON-RAMOS 2009; YOSHII AND CONSTANTINE-PATON 2010; CHOU *et al*. 2020; BHARMAURIA *et al*. 2022; HEDRICK *et al*. 2024). Disruptions in these pathways are implicated in various neurodevelopmental and neurodegenerative disorders (WASHBOURNE 2015; DEJANOVIC *et al*. 2024). Consequently, these structural changes are tightly controlled by intracellular signaling networks and membrane trafficking pathways that coordinate synaptic growth and maintenance (CRAVEN AND BREDT 1998; DESHPANDE AND RODAL 2016; CHOU *et al*. 2020; JIANG *et al*. 2021). While several core regulators of synaptic growth have been identified, the molecular mechanisms that fine-tune neuronal signaling and integrate them with intracellular trafficking remain incompletely understood.

Endocytosis, particularly Clathrin-Mediated Endocytosis (CME), plays a central role in synaptic development by regulating receptor trafficking, signal attenuation, and synaptic vesicle recycling (DI FIORE AND DE CAMILLI 2001; ROYLE AND LAGNADO 2003; HEERSSEN *et al*. 2008). For instance, mutations in endocytic components, including Dynamin, Nervous Wreck (Nwk), AP2 subunits, Synaptojanin 1 (Synj1), or Endophilin (Endo), lead to aberrant synaptic growth, characterized by excessive bouton number, altered bouton size, and/or disrupted active zone spacing (DICKMAN *et al*. 2006; O’CONNOR-GILES *et al*. 2008; ZHAO *et al*. 2013; LAUGKS *et al*. 2017). These phenotypes are primarily attributed to hyperactivation of the BMP signaling pathway (KESHISHIAN AND KIM 2004; MCCABE *et al*. 2004). The BMP pathway, activated by muscle-derived Glass bottom boat (Gbb) binding to Thickveins/Saxophone/Wishful thinking (Tkv/Sax/Wit) receptors, induces Smad-mediated gene expression that promotes synaptic growth (MCCABE *et al*. 2003; GOOLD AND DAVIS 2007; BALL *et al*. 2010; RODAL *et al*. 2011; SMITH *et al*. 2012; VANLANDINGHAM *et al*. 2013). While endocytosis modulates this signaling by regulating receptor trafficking, and its disruption enhances BMP activity (DICKMAN *et al*. 2006; CHOUDHURY *et al*. 2022), the BMP-independent roles of endocytosis in synapse growth signaling remain largely unknown.

Genetic studies in *Drosophila* indicate that suppressing BMP signaling alone does not fully rescue the synaptic overgrowth observed in endocytic mutants (COLLINS AND DIANTONIO 2007a; CHOUDHURY *et al*. 2022), suggesting that additional, parallel pathways are required to restore normal synaptic morphology. One such pathway is the Wnd/DLK-JNK cascade, which may act as an additional signaling pathway for synaptic remodeling. Consistent with such a hypothesis, Hiw/Phr1 links autophagy and MAP3K signaling by targeting Wnd for degradation (COLLINS *et al*. 2006; WU *et al*. 2007). While autophagy induces synaptic growth by reducing Hiw levels (SHEN AND GANETZKY 2009), Hiw negatively regulates synaptic growth by targeting Wnd, a key activator of the JNK pathway, for degradation (COLLINS *et al*. 2006; WU *et al*. 2007). Loss of Hiw leads to persistent JNK activation and NMJ overgrowth, and is similar to defects in endocytic mutants. Hiw also promotes the degradation of co-Smad Medea, implicating it as a key regulator of both BMP and JNK pathways (MCCABE *et al*. 2004). While mutations in Medea only partially suppress the synaptic overgrowth, *wnd* mutants completely suppress the synaptic overgrowth observed in the *hiw* mutants (COLLINS *et al*. 2006; COLLINS AND DIANTONIO 2007a). These genetic epistatic studies imply that, similar to *hiw* mutants, endocytic mutants might regulate multiple signaling pathways that restrict synaptic growth.

Here, we identify a previously unrecognized role for endocytic proteins in restricting synaptic growth by regulating the trafficking, localization, and activity of Hiw. Our study reveals that impaired endocytosis leads to the accumulation and mislocalization of Hiw into dense, dynamic punctate structures within neuronal cell bodies. These aberrant Hiw assemblies cannot target Wnd for degradation, resulting in elevated Wnd levels and sustained activation of JNK signaling. We further demonstrate that Rab11-positive recycling endosomes are crucial for Hiw trafficking, as Rab11 loss recapitulates endocytic mutant phenotypes, including Hiw puncta formation and Wnd upregulation. Our findings support a model in which endocytosis, via Rab11-mediated recycling and autophagy, possibly ensures proper localization and turnover of Hiw/Phr1, preventing its sequestration into nonfunctional protein assemblies and thereby limiting Wnd-JNK signaling. Thus, our work expands the functional repertoire of endocytic proteins and underscores their essential role in maintaining synaptic homeostasis.

## Results

### Endocytic Impairment Drives Hiw/Phr1 Accumulation and Synaptic Overgrowth Independent of Autophagy

Endocytosis is an established regulator of *Drosophila* NMJ growth, in part because defective endocytic machinery elevates retrograde BMP signaling, leading to synaptic overgrowth (O’CONNOR-GILES *et al*. 2008; BALL *et al*. 2010; RODAL *et al*. 2011; CHOUDHURY *et al*. 2022). However, blocking BMP signaling only partially suppresses the overgrowth observed in several endocytic mutants (COLLINS AND DIANTONIO 2007b; CHOUDHURY *et al*. 2022), indicating that additional pathways contribute to this phenotype. This motivated us to investigate whether altered Hiw-dependent JNK signaling contributes to this process. Hiw is one of the key regulators of JNK signaling and synaptic growth. It negatively regulates synaptic growth by targeting the MAP3K, Wnd, for ubiquitin-proteasomal degradation (WAN *et al*. 2000; COLLINS *et al*. 2006).

Due to the lack of a reliable antibody against Hiw, we used GFP-tagged Hiw to analyze Hiw/Phr1 levels in the endocytic mutants. The GFP-Hiw transgene has been extensively used to analyze the functional roles of Hiw in *Drosophila* synaptic growth (WU *et al*. 2005a; SHEN AND GANETZKY 2009), and learning and memory (SHIN AND DIANTONIO 2011). Neuronal expression of GFP-Hiw in the background of endocytic mutants showed a significant increase in Hiw levels in the third-instar larval ventral nerve cord (VNC) (Figure 1A-I and 1L-M). Careful examination of Hiw organization in these mutants revealed formation of large cytoplasmic puncta in motor neuron cell bodies (Figure 1C-K). While the expression of GFP-Hiw transgenes in an otherwise wild-type background does not cause significant Hiw accumulation, the large punctate accumulations were seen only when expressed in the endocytic mutants (Figure 1J-K). Importantly, the expression of GFP alone in *AP2* mutants did not induce GFP accumulations or increased GFP intensity, confirming that the phenotype was not due to overexpression artifacts (Figure 1B-K). Consistently, knockdown of other AP2 subunits (AP2α/AP2β) also induced the formation of Hiw puncta in neuronal cell bodies (Figure 1-figure supplement 1 A-D). Hiw is known to traffic to synapses and regulate target protein levels, such as Wnd and NMNAT, thereby regulating synaptic growth and axon regeneration pathways, respectively (XIONG *et al*. 2012a; RUSSO *et al*. 2019). Concurrently, our analysis of Hiw levels at the NMJ revealed a significant reduction in Hiw levels at the NMJ in *AP2* or *synj1* mutants (Figure 1-figure supplement 1 E-I), suggesting that Hiw accumulation in cell bodies likely impairs its trafficking to synaptic terminals.

**Figure 1:**
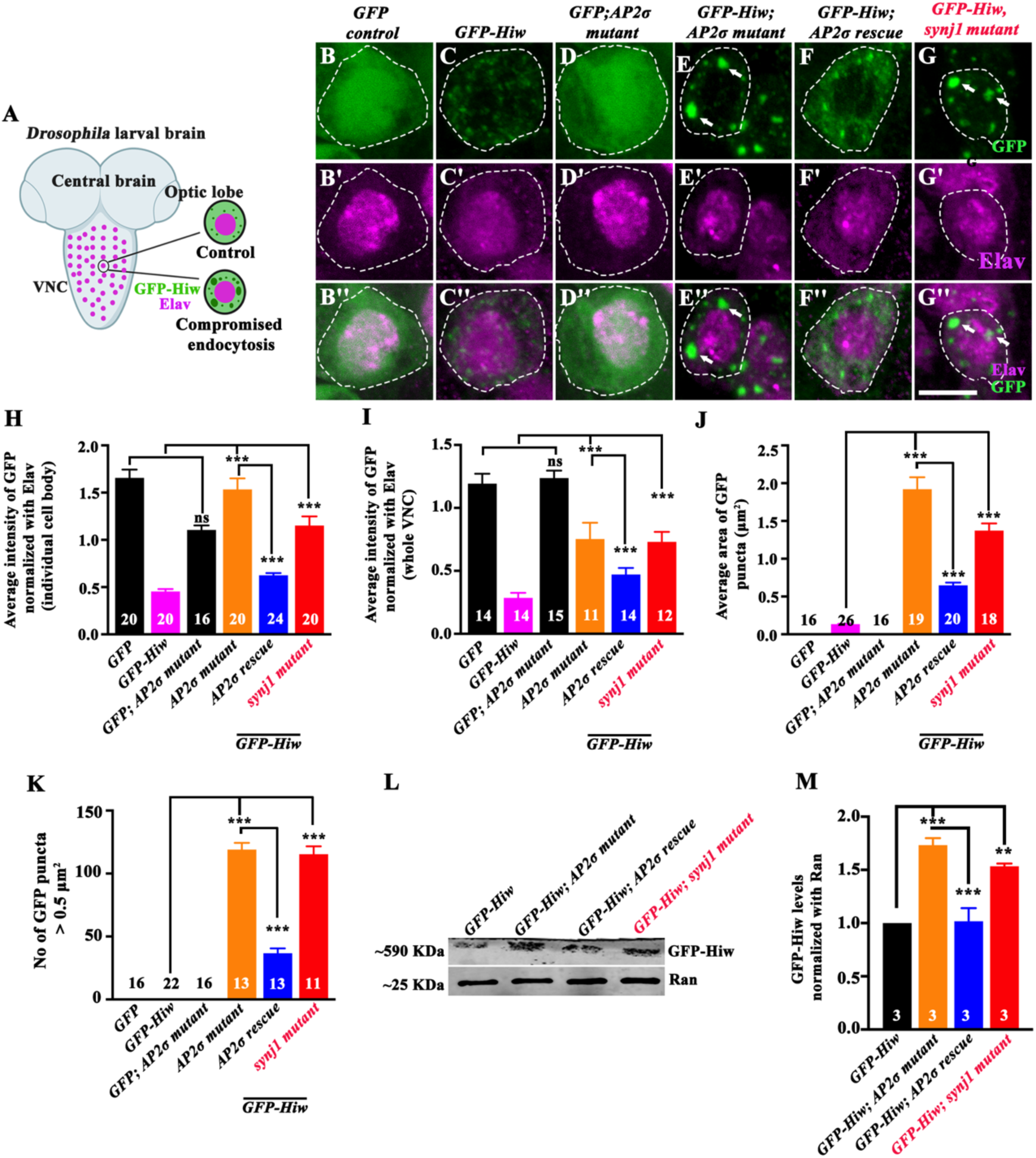
Compromised endocytosis causes Hiw to accumulate in large punctae within the neuronal cell body. (**A**) Schematic representation of the third instar larval ventral nerve cord (VNC). The magnified view shows accumulation and possible mislocalization of Hiw assemblies in neuronal cell bodies of endocytic mutants. (**B-G’’**) Confocal images of the neuronal cell body in third instar larval VNC co-labeled for GFP-Hiw (green) and Elav (magenta) in GFP control (*Elav-Gal4>UAS-GFP)* (B-B’’)*, Elav-Gal4>UAS-GFP-Hiw* (C-C’’), *Elav-Gal4>UAS-GFP; AP2σ-^KG^/AP2σ^ang7^* (D-D’’), *Elav-Gal4>UAS-GFP-Hiw; AP2σ-^KG^/AP2σ^ang7^* (E-E’’), *Elav-Gal4>UAS-GFP-Hiw; UAS-AP2σ, AP2σ-^KG^/AP2σ^ang7^* (F-F’’), and *Elav-Gal4>UAS-GFP-Hiw; synj1^1/2^* (G-G’’). The scale bar in G’’ represents 5 µm. (**H**) Histogram showing the average intensity of GFP in individual neuronal cell body in GFP control (*Elav-Gal4>UAS-GFP*) (1.66 ± 0.08)*, Elav-Gal4>UAS-GFP-Hiw* (0.45 ± 0.02), *Elav-Gal4>UAS-GFP; AP2σ-^KG^/AP2σ^ang7^*(1.11 ± 0.04), *Elav-Gal4>UAS-GFP-Hiw; AP2σ-^KG^/AP2σ^ang7^* (1.53 ± 0.11), *Elav-Gal4>UAS-GFP-Hiw; UAS-AP2σ, AP2σ-^KG^/AP2σ^ang7^* (0.62 ± 0.02), and *Elav-Gal4>UAS-GFP-Hiw; synj1^1/2^* (1.15 ± 0.09). At least 16 VNCs from each genotype were used for quantification. (I) Histogram showing the average intensity of GFP-Hiw in third instar larval VNC in GFP control (*Elav-Gal4>UAS-GFP*) (1.19 ± 0.07)*, Elav-Gal4>UAS-GFP-Hiw* (0.28 ± 0.04), *Elav-Gal4>UAS-GFP; AP2σ-^KG^/AP2σ^ang7^*(1.24 ± 0.05), *Elav-Gal4>UAS-GFP-Hiw; AP2σ-^KG^/AP2σ^ang7^* (0.753 ± 0.12), *Elav-Gal4>UAS-GFP-Hiw; UAS-AP2σ, AP2σ-^KG^/AP2σ^ang7^* (0.471 ± 0.05), and *Elav-Gal4>UAS-GFP-Hiw; synj1^1/2^*(0.73 ± 0.07). At least 12 VNCs from each genotype were used for quantification. (J) Histogram showing the average area (μm^2^) of GFP-Hiw puncta in GFP control (*Elav-Gal4>UAS-GFP*) (0.00 ± 0.00)*, Elav-Gal4>UAS-GFP-Hiw* (0.13 ± 0.01), *Elav-Gal4>UAS-GFP; AP2σ-^KG^/AP2σ^ang7^* (0.00 ± 0.00), *Elav-Gal4>UAS-GFP-Hiw; AP2σ-^KG^/AP2σ^ang7^* (1.92 ± 0.15), *Elav-Gal4>UAS-GFP-Hiw; UAS-AP2σ, AP2σ-^KG^/AP2σ^ang7^* (0.64 ± 0.03), and *Elav-Gal4>UAS-GFP-Hiw; synj1^1/2^* (1.37 ± 0.09). At least 16 VNCs from each genotype were used for quantification. (K) Histogram showing the number of GFP puncta with an area greater than 0.5 μm^2^ in GFP control (*Elav-Gal4>UAS-GFP*) (0.00 ± 0.00)*, Elav-Gal4>UAS-GFP-Hiw* (0.00 ± 0.00), *Elav-Gal4>UAS-GFP; AP2σ-^KG^/AP2σ^ang7^*(0.00 ± 0.00), *Elav-Gal4>UAS-GFP-Hiw; AP2σ-^KG^/AP2σ^ang7^* (119 ± 5.35), *Elav-Gal4>UAS-GFP-Hiw; UAS-AP2σ, AP2σ-^KG^/AP2σ^ang7^* (36.7 ± 3.87), and *Elav-Gal4>UAS-GFP-Hiw; synj1^1/2^* (115 ± 6.26). At least 11 VNCs from each genotype were used for quantification. (L) Western blot showing the level of GFP-Hiw in control (*Elav-Gal4>UAS-GFP-Hiw*), *Elav-Gal4>UAS-GFP-Hiw; AP2σ-^KG^/AP2σ^ang7^*, *Elav-Gal4>UAS-GFP-Hiw; UAS-AP2σ, AP2σ-^KG^/AP2σ^ang7,^* and *Elav-Gal4>UAS-GFP-Hiw; synj1^1/2^*. Ran was used as a loading control. (M) Histogram showing the level of GFP-Hiw in control (*Elav-Gal4>UAS-GFP-Hiw*) (1.0 ± 0.00), *Elav-Gal4>UAS-GFP-Hiw; AP2σ-^KG^/AP2σ^ang7^* (1.73 ± 0.06), *Elav-Gal4>UAS-GFP-Hiw; UAS-AP2σ, AP2σ-^KG^/AP2σ^ang7^* (1.02 ± 0.12), and *Elav-Gal4>UAS-GFP-Hiw; synj1^1/2^* (1.53 ± 0.02). Three independent Western blots were used for quantification. The error bars in H, I, J, K, and M represent the standard error of the mean (SEM); the statistical analysis was performed using one-way ANOVA followed by Tukey’s post hoc test. ***p<0.001; **p<0.01 ns: not significant.

Autophagy serves as an additional layer of regulation for Hiw and thereby modulates synaptic growth. Under normal conditions, Hiw protein levels are regulated through selective autophagic degradation (SHEN AND GANETZKY 2009). Our observation of increased Hiw levels and their accumulation in the endocytic mutants, together with the well-established functional interplay between endocytosis and autophagy (TOOZE *et al*. 2014; KAKUTA *et al*. 2017; BIRGISDOTTIR AND JOHANSEN 2020), and the role of CME in autophagosome biogenesis (TIAN *et al*. 2013; SOUKUP *et al*. 2016; KONONENKO *et al*. 2017; HERNANDEZ-DIAZ *et al*. 2022), prompted us to investigate whether autophagy has any role in mediating the synaptic overgrowth in the endocytic mutants. To test this, we first examined the autophagy status in *Drosophila AP2* mutants by assessing Atg8a levels at the NMJ using an Atg8a-GFP reporter (LOW *et al*. 2013; MAUVEZIN *et al*. 2014). We observed a reduction in Atg8a-GFP levels in *AP2* mutant neurons, suggesting that autophagy might be compromised in these mutants (Figure 1 – figure supplement 2). To further assess autophagic flux, we employed the tandem-tagged GFP-mCherry-Atg8a reporter, a well-established marker for assessing autophagic flux *in vivo* (Figure 1 – figure supplement 3A) (DEVORKIN AND GORSKI 2014). Consistently, we found decreased fluorescence of both GFP and mCherry signals in *AP2* mutants (Figure 1 – figure supplement 3B-H). Similar results were obtained in *synj1* mutants (Figure 1 – figure supplement 3E-H), indicating that although autophagic flux *per se* may not be severely impaired, autophagosome formation is likely compromised in endocytic mutants, thereby reducing autophagy. Next, to determine whether defective autophagy contributes to the synaptic overgrowth phenotype. Autophagy positively regulates synaptic overgrowth, as autophagy mutants exhibit reduced synaptic growth due to elevated Hiw levels (SHEN AND GANETZKY 2009). Similarly, we genetically manipulated autophagy in *AP2* and *synj1* mutant backgrounds (*atg1^Δ3D^/atg1^Δ3D^, AP2σ^KG/Ang7^* or *synj1^1/2^; atg1^Δ3D^/atg1^Δ3D^*). The double mutants showed synaptic overgrowth similar to the *AP2* and *synj1* mutants (Figure 1 - figure Supplement 4), indicating that although endocytic defects perturb autophagy, these pertubations alone do not account for the observed synaptic overgrowth. These findings suggest that disruption of autophagy may not be the primary cause of synaptic phenotypes observed in the endocytic mutants.

### Endocytic Disruption Promotes the Accumulation of Dynamic Hiw/Phr1 Puncta

Given that compromised endocytosis led to the accumulation of Hiw/Phr1 in large cytoplasmic puncta, we hypothesized that Hiw may possess intrinsic biophysical features that enable formation of assemblies, and that these assemblies might correspond to biomolecular condensates rather than aggregates. To test this, we performed a comparative *in silico* analysis using Prion-Like Amino Acid Composition (PLAAC) (LANCASTER *et al*. 2014), IUPred2A, and ANCHOR2 (MESZAROS *et al*. 2018), with Ataxin-2 (Atx2) and GFP serving as positive and negative controls, respectively (BAKTHAVACHALU *et al*. 2018) (Figure 2A-B). As expected, GFP exhibited minimal disorder or binding propensity (IUPred2A/ANCHOR2 scores < 0.4 throughout) and a negative log-likelihood ratio (LLR = -9.614) in PLAAC, suggesting a low propensity for condensation. In contrast, Atx2 showed widespread highly disordered regions, with both IUPred2A and ANCHOR2 scores exceeding the threshold across most of the sequence. PLAAC analysis of Atx2 yielded a strong LLR score of 45.352, in line with its known role in RNP granule formation and stress-induced phase separation (BECKER *et al*. 2017; BAKTHAVACHALU *et al*. 2018) (Figure 2A-B).

**Figure 2:**
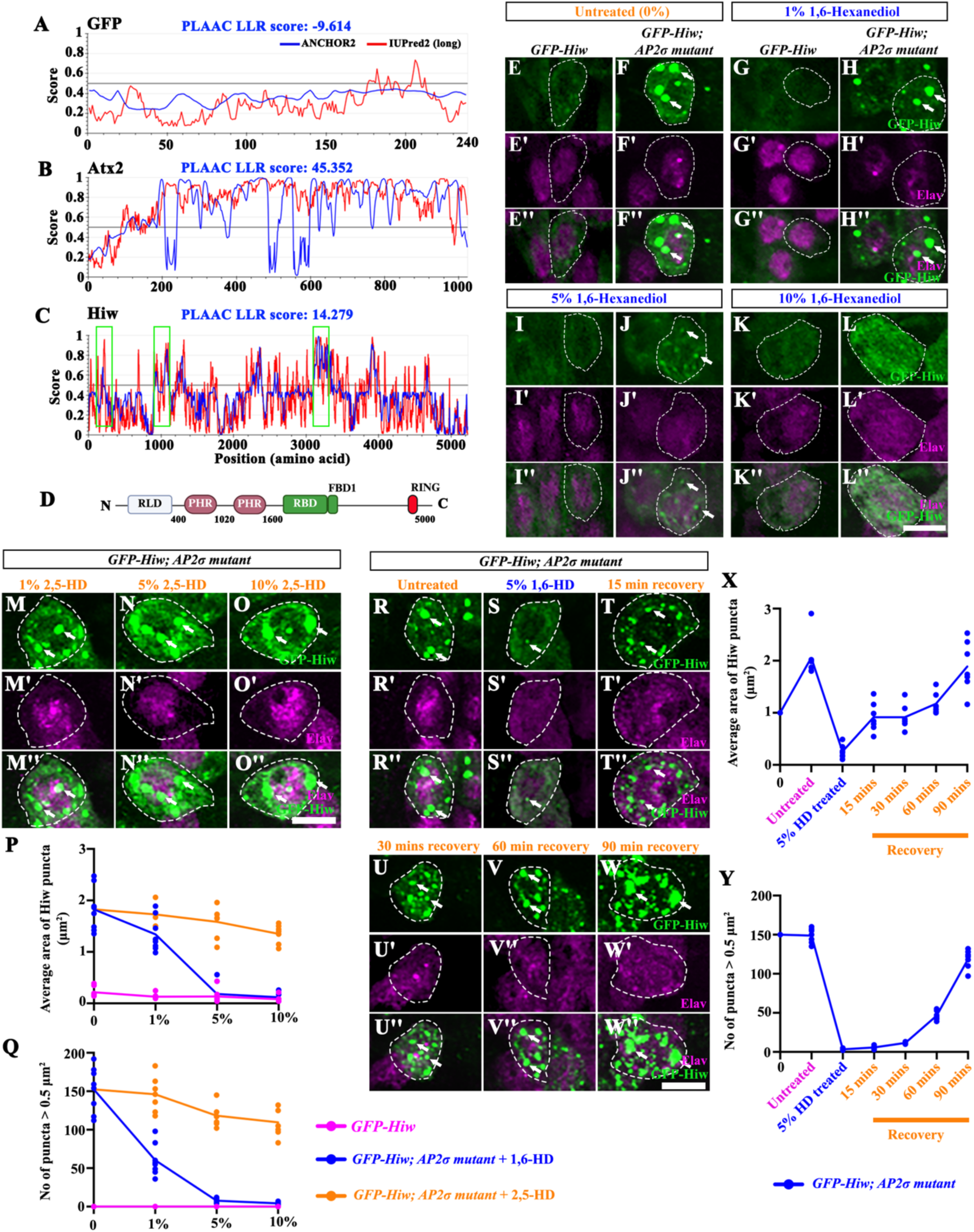
Highwire punctate assemblies are dynamic in endocytic mutants. (**A-C**) The PLAAC scores of GFP, Atx2, and Hiw. The PLAAC scores represent the ability of a protein segment to form aggregates or LLPS (LANCASTER *et al*. 2014). The plots represent predicted intrinsically disordered regions in a protein. The red line denotes the likelihood of a protein segment being disordered. The blue line denotes the probability of a given position in a protein being a part of a disordered binding region. (**D**) The domain organization of the Hiw protein. It contains an N-terminal RCC1-like domain (RLD), two (PAM-Hiw-RPM-1) PHR domains, followed by a (Rae1-binding domain) RBD and a (Fsn-1 binding domain) FBD1, and a C-terminal (RING H2 ubiquitin-ligase domain) RING domain. (**E-L’’**) Confocal images of the neuronal cell body co-labelled for GFP-Hiw (green) and Elav (magenta) in control (*Elav-Gal4>UAS-GFP-Hiw*) and *Elav-Gal4>UAS-GFP-Hiw; AP2σ-^KG^/AP2σ^ang7^* treated with 0% 1,6-HD (E-F’’), 1% 1,6-HD (G-H’’), 5% 1,6-HD (I-J’’), and 10% 1,6-HD (K-L’’) containing Schneider’s media for 2 mins. The 0% contains only media. The scale bar in L’’ represents 5 µm for images E-L’’. (**M-O’’**) Confocal images of the neuronal cell body co-labeled for GFP-Hiw (green) and Elav (magenta) in *Elav-Gal4>UAS-GFP-Hiw; AP2σ-^KG^/AP2σ^ang7^* treated with 1% 2,5-HD (M-M’’), 5% 2,5-HD (N-N’’), and 10% 2,5-HD (O-O’’) containing Schneider’s medium for 2 mins. The scale bar in O’’ represents 5 µm for images M-O’’. (**P**) Line graph showing the average area (μm^2^) of the GFP-Hiw punctae in control (*Elav-Gal4>UAS-GFP-Hiw*) (magenta line) and *Elav-Gal4>UAS-GFP-Hiw; AP2σ^KG^/AP2σ^ang7^* treated with 0%, 1%, 5% and 10% 1,6-HD (blue line), and *Elav-Gal4>UAS-GFP-Hiw; AP2σ^KG^/AP2σ^ang7^* treated with 0%, 1%, 5% and 10% 2,5-HD (orange line). (**Q**) Line graph showing the number of the GFP-Hiw punctae with an area greater than 0.5 μm^2^ in control (*Elav-Gal4>UAS-GFP-Hiw*) (magenta line) and *Elav-Gal4>UAS-GFP-Hiw; AP2σ^KG^/AP2σ^ang7^* treated with 0%, 1%, 5% and 10% 1,6-HD (blue line), and *Elav-Gal4>UAS-GFP-Hiw; AP2σ^KG^/AP2σ^ang7^* treated with 0%, 1%, 5% and 10% 2,5-HD (orange line). (**R-W’’**) Confocal images of the neuronal cell body co-labelled for GFP-Hiw (green) and Elav (magenta) in *Elav-Gal4>UAS-GFP-Hiw; AP2σ-^KG^/AP2σ^ang7^* ( R-R’’) treated with 5% 1,6-HD for 2 min (S-S’’) and allowed to recover for 15 mins (T-T’’), 30 mins (U-U’’), 60 mins (V-V’’), and 90 mins (W-W’’) after removal of 1,6-HD in S2 Schneider’s media. The scale bar in W’’ represents 5 µm. (**X**) Line graph showing the area (µm^2^) of GFP-Hiw puncta in *Elav-Gal4>UAS-GFP-Hiw; AP2σ-^KG^/AP2σ^ang7^* treated with 5% 1,6-HD for 2 min and after recovery for 15 mins, 30 mins, 60 mins, and 90 mins. At least 8 VNCs from each genotype were used for quantification. (**Y**) Line graph showing the number of GFP-Hiw puncta with an area greater than 0.5 µm^2^ in *Elav-Gal4>UAS-GFP-Hiw; AP2σ-^KG^/AP2σ^ang7^* treated with 5% 1,6-HD for 2 min, and after recovery for 15 mins, 30 mins, 60 mins, and 90 mins. At least 8 VNCs from each genotype were used for quantification.

Remarkably, Hiw displayed intermediate but significant aggregation potential. PLAAC analysis identified several Q/N-rich prion-like regions, with an LLR of 14.279, suggesting bona fide prion-like content. IUPred2A and ANCHOR2 predictions showed distinct stretches of high disorder and disordered binding potential, notably within the N-terminal RCC1-like region (residues ∼200–400), a central segment (∼900-1050), and an extended region between residues 3000-3700 (Figure 2C-D). Compared to Atx2, Hiw harbors fewer but sharper peaks of disorder and interaction propensity, suggesting a modular architecture capable of regulated condensate formation.

To investigate the nature of Hiw puncta in *AP2* mutants, we examined colocalization with endosomal and stress granule markers. We found that Hiw puncta do not colocalize with markers of early (Rab5), late (Rab7), or recycling (Rab11) endosomes (Figure 2 - figure supplement 1A-F’’), suggesting that it might not be part of these classical vesicular structures. Similarly, we did not find colocalization with the stress granule marker, Caprin, or the integrated stress response marker, p-eIF2α (Figure 2 - figure supplement 1G-J’’).

Given their punctate appearance and lack of colocalization with the endosomal markers, we asked whether these structures were sensitive to disruption of weak molecular interactions. To address this, we next tested whether Hiw puncta were phase-separated condensates or stable aggregates. Treatment with 1,6-hexanediol (1,6-HD), which disrupts weak hydrophobic interactions in liquid-like assemblies (KROSCHWALD *et al*. 2017), caused a rapid, concentration-dependent dissolution of GFP-Hiw puncta (Figure 2E-L and 2P-Q). In contrast, treatment with 2,5-hexanediol, a similar aliphatic alcohol (NAIR *et al*. 2019), did not dissolve the Hiw/Phr1 puncta, strongly supporting that Hiw/Phr1 puncta were formed by non-covalent, likely hydrophobic interactions between Hiw (Figure 2M-Q). Importantly, the assembly of Hiw-GFP puncta was reversible, as washing out 1,6-HD and incubating the brains in Schneider’s medium reassembled the Hiw puncta in a time-dependent manner, indicating that the Hiw/Phr1 puncta are maintained by dynamic, reversible molecular interactions (Figure 2R-Y). Further, we attempted fluorescence recovery after photobleaching (FRAP) on these Hiw accumulations. However, due to the large size of the GFP-Hiw protein (∼590 kDa) and the limitations of *in vivo* imaging in the larval VNC, we were unable to reliably detect a significant recovery of Hiw following photobleaching of the Hiw puncta. Together, these data support the view that the observed Hiw/Phr1 puncta are dynamic and maintained by weak, reversible interactions rather than insoluble aggregates. However, their definitive biophysical nature requires further investigations.

To further distinguish whether the Hiw accumulation in the cell body reflected an endosomal trafficking or the autophagy-dependent degradation defect, we examined Hiw distribution in pharmacologically induced (using Bafilomycin A1 or Chloroquine) or genetically (*atg1^Δ3D^*) autophagy-deficient animals (SCOTT *et al*. 2004). We found that although overall GFP-Hiw fluorescence increased in autophagy-deficient animals, large cytoplasmic puncta were not observed (Figure 2 - figure supplement 2), indicating that endosomal trafficking disruption, and not degradation, is likely responsible for Hiw accumulation in endocytic mutants.

Together, these results support that: a) the puncta observed in the endocytic mutants represent dynamic Hiw assemblies; and b) the Hiw puncta formation is independent of autophagy.

### Acute Endocytic Block Triggers Formation of Hiw/Phr1 Assemblies

Since the *AP2* and *synj1* mutants we analyzed were loss-of-function alleles with impaired endocytosis throughout development, we next asked whether acute inhibition of endocytosis was sufficient to induce Hiw/Phr1 puncta formation. To test this, we examined temperature-sensitive *Shibire* (*shi^ts^*) mutants, which harbor dynamin mutations that cause a reversible block in endocytosis at restrictive temperatures (KOSAKA AND IKEDA 1983). Hiw-positive puncta were detectable after 3 hours at the non-permissive temperature, and both puncta size and number increased following 6 hours of endocytic inhibition. These observations indicate that acute endocytic inhibition is associated with the time-dependent formation and accumulation of Hiw puncta in neuronal cell bodies (Figure 3B-K’ and 3M-N). These assemblies resembled those seen in *AP2* or *synj1* mutants and were absent under permissive conditions (Figure 3A-D’ and 3M-N), suggesting that Hiw/Phr1 assemblies arise due to endocytic blockade rather than developmental effects.

**Figure 3:**
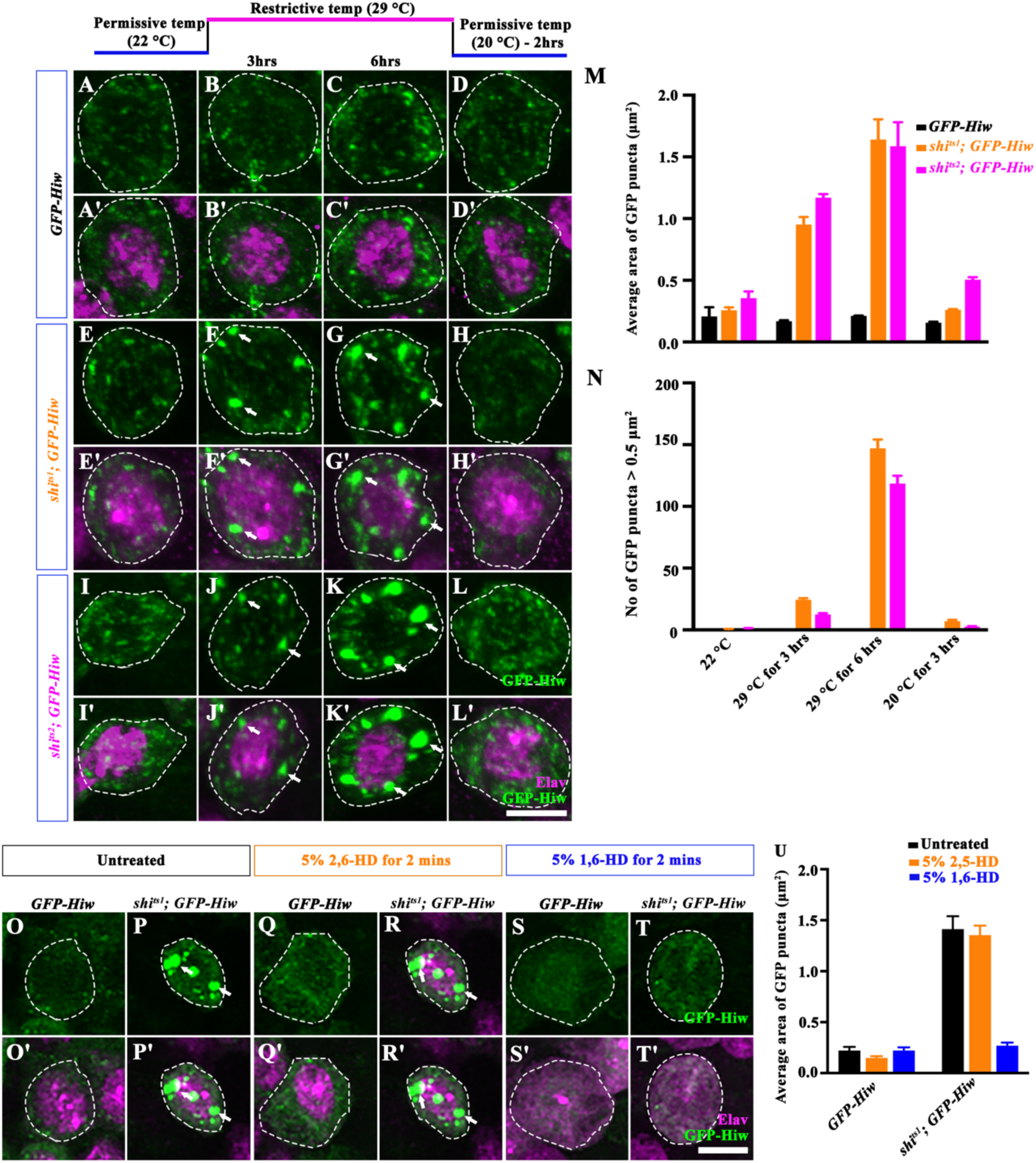
Acute inhibition of endocytosis triggers Hiw/Phr1 accumulation. (**A-L’**) Confocal images of the neuronal cell body co-labeled for GFP-Hiw (green) and Elav (magenta) in control (*Elav-Gal4>UAS-GFP-Hiw*) (A-D’), *Elav-Gal4>UAS-GFP-Hiw; shi^ts1^* (E-H’), and *Elav-Gal4>UAS-GFP-Hiw; shi^ts2^*(I-L’). The scale bar in L’ represents 5 µm. (**M**) Histogram showing the average area (µm^2^) of GFP-Hiw puncta in control, *Elav-Gal4>UAS-GFP-Hiw; shi^ts1^*, and *Elav-Gal4>UAS-GFP-Hiw; shi^ts2^* at permissive (22 °C) and restrictive temperatures (29 °C) at 3 hrs and 6 hrs, followed by recovery at permissive temperature (20 °C) for 2 hrs. At least 8 VNCs from each genotype were used for quantification. (**N**) Histogram showing the number of GFP-Hiw puncta with an area greater than 0.5 µm^2^ in control, *Elav-Gal4>UAS-GFP-Hiw; shi^ts1^*, and *Elav-Gal4>UAS-GFP-Hiw; shi^ts2^* at permissive (22 °C) and restrictive temperatures (29 °C) at 3 hrs and 6 hrs, followed by recovery at permissive temperature (20 °C) for 2 hrs. At least 8 VNCs from each genotype were used for quantification. The error bar in M and N represents the standard error of the mean (SEM). (**O-T’**) Confocal images of the neuronal cell body co-labeled for GFP-Hiw (green) and Elav (magenta) in *Elav-Gal4>UAS-GFP-Hiw and Elav-Gal4>UAS-GFP-Hiw; shi^ts1^* treated with 5% 2,5-HD and 1,6-HD for 2 min. The scale bar in T’ represents 5 µm. (**U**) Histogram showing the average area of GFP-Hiw puncta in untreated, 5% 2,5-HD, and 5% 1,6-HD treated controls, and *Elav-Gal4>UAS-GFP-Hiw; shi^ts1^*. At least 8 VNCs from each genotype were used for quantification. The error bar represents the standard error of the mean (SEM).

We next asked whether Hiw/Phr1 assemblies formed in response to an acute endocytic block were dynamic. We found that the Hiw/Phr1 puncta induced by acute endocytic blockade disassemble with a temperature shift from restrictive to permissive conditions (Figure 3H-L’ and 3M-N). Moreover, the puncta induced at restrictive temperature were sensitive to 1,6-HD but not to 2,5-HD, supporting the conclusion that the observed Hiw/Phr1 puncta are dynamic in nature (Figure 3O-U). Together, these results confirm that endocytic impairments sequester Hiw/Phr1 into dynamic and reversible protein accumulations.

### Wnd/DLK Upregulation Drives the Synaptic Overgrowth in Endocytic Mutants

Hiw accumulates in neuronal cell bodies following disruption of endocytosis, with reduced synaptic Hiw levels. We therefore asked whether the accumulated Hiw retains its ability to restrain synaptic growth or whether accumulation in the cell body compromises its functional activity. To address this, we expressed *GFP-Hiw* in *AP2* mutants (*Elav-Gal4>UAS-GFP-Hiw; AP2σ^KG/Ang7^*) and assessed whether it could suppress the synaptic overgrowth phenotype. We found that GFP-Hiw expression in endocytic mutants failed to rescue synaptic overgrowth (Figure 4 - figure supplement 1), suggesting that increasing Hiw expression is insufficient to restore its growth-restraining function when endocytosis is impaired. This finding is consistent with the possibility that altered Hiw localization or reduced synaptic availability contributes to the compromised functions of Hiw in endocytic mutants.

Wnd promotes synaptic growth at the *Drosophila* NMJ by activating the JNK signaling pathway. Hiw restrains this growth mainly by targeting Wnd for degradation. Since accumulated Hiw in the *AP2* mutant failed to suppress synaptic overgrowth and was functionally compromised, we asked whether Wnd levels were altered in endocytic mutants. We found that *AP2* mutants exhibited elevated Wnd levels in the VNCs, which were restored to control levels upon neuronal expression of an *AP2* transgene (Figure 4A-F). Similarly, *synj1* mutants also displayed both Hiw accumulation and increased Wnd levels in neurons (Figure 4E-F). Consistent with these findings, we found that endocytic mutants showed elevated levels of an endogenously tagged mCherry-Wnd (VENKEN *et al*. 2011; KIM *et al*. 2024) (Figure G-J).

**Figure 4:**
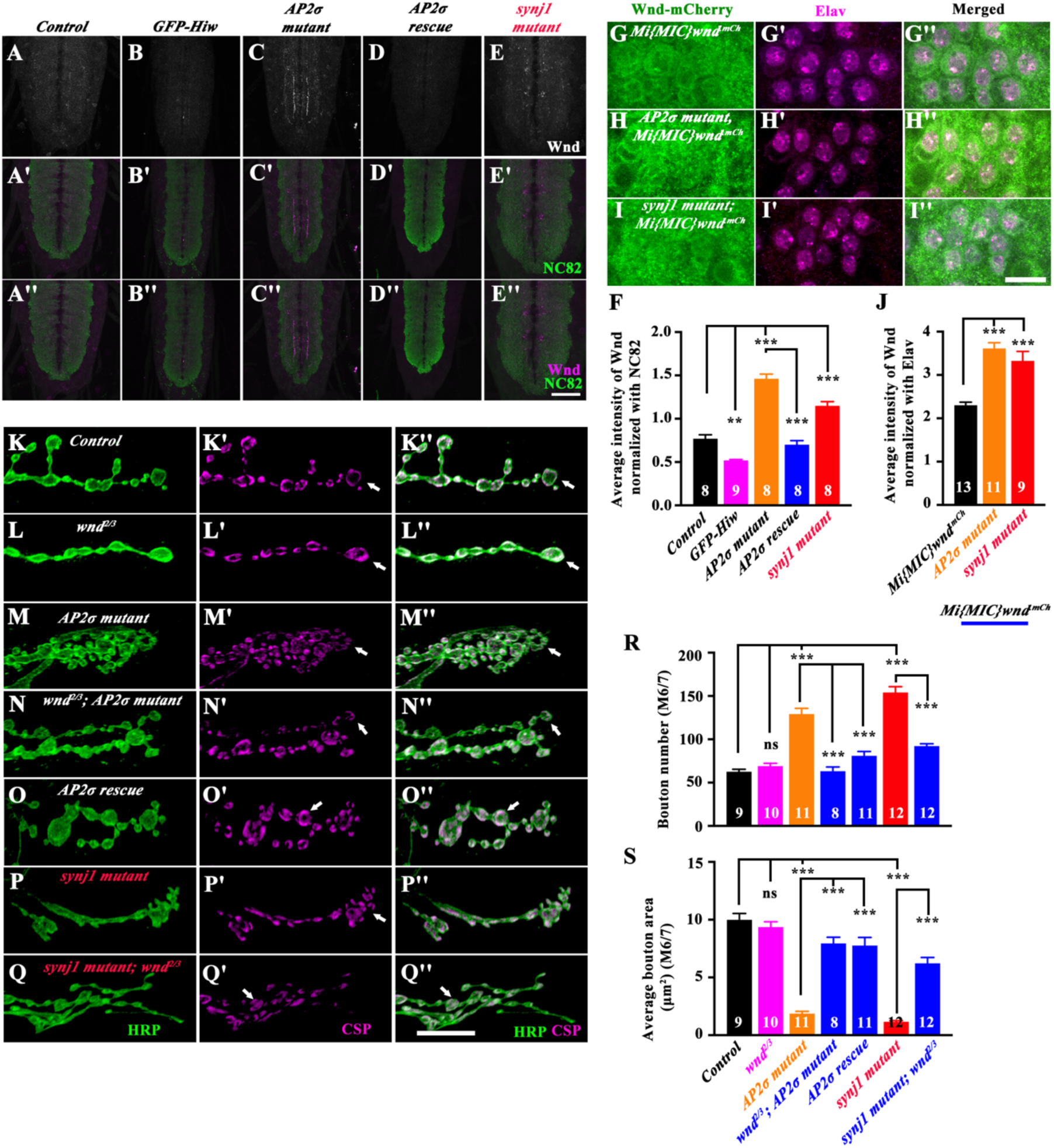
Endocytosis restricts synaptic growth via MAP3K/Wnd signaling. (**A-E’’**) Confocal images of third instar larval ventral nerve cord co-labeled for Wnd (magenta) and NC82 (green) in control (A-A’’), *Elav-Gal4>UAS-GFP-Hiw* (B-B’’), *AP2σ-^KG^/AP2σ^ang7^* (C-C’’), *Elav-Gal4> UAS-AP2σ, AP2σ-^KG^/AP2σ^ang7^* (D-D’’), and *synj1^1/2^* (E-E’’). The scale bar in E’’ represents 10 µm. (**F**) Histogram showing the average fluorescence intensity of Wnd normalized with NC82 in control (0.77 ± 0.04), *Elav-Gal4>UAS-GFP-Hiw* (0.52 ± 0.01), *AP2σ-^KG^/AP2σ^ang7^* ( 1.46 ± 0.05), *Elav-Gal4> UAS-AP2σ, AP2σ-^KG^/AP2σ^ang7^* (0.71± 0.04) and *synj1^1/2^*; (1.15 ± 0.04). At least 8 VNCs of each genotype were used for quantification. The error bar represents the standard error of the mean (SEM); the statistical analysis was done using one-way ANOVA followed by post hoc Tukey’s test. ***p<0.001; **p<0.01. (**G-I’’**) Confocal images of third-instar larval ventral nerve cord co-labeled for Wnd (green) and Elav (magenta) in control (*Mi{MIC}wnd^.mCh^)* (G-G’’), *AP2σ-^KG^/AP2σ^ang7^; Mi{MIC}wnd^.mCh^* (H-H’’), and *synj1^1/2^*; *Mi{MIC}wnd^.mCh^* (I-I’’). The scale bar in I’’ represents 10 µm. (**J**) Histogram showing the average fluorescence intensity of Wnd in control (Endogenously tagged Wnd-mCherry; *Mi{MIC}wnd^.mCh^*) (2.30 ± 0.07), *AP2σ-^KG^/AP2σ^ang7^*; *Mi{MIC}wnd^.mCh^* ( 3.61 ± 0.13), and *synj1^1/2^*; *Mi{MIC}wnd^.mCh^* (3.33 ± 0.21). At least 9 VNCs of each genotype were used for quantification. The error bar represents the standard error of the mean (SEM); the statistical analysis was done using one-way ANOVA followed by post hoc Tukey’s test. ***p<0.001. (**K-Q’’**) Confocal images of muscle 6/7 NMJ at A2 hemisegment showing synaptic growth in control (K-K’’), *wnd^2/3^* (L-L’’)*, AP2σ-^KG^/AP2σ^ang7^*(M-M’’)*, wnd^2/3^, AP2σ-^KG^/AP2σ^ang7^* (N-N’’), *Elav-Gal4>UAS-AP2σ, AP2σ-^KG^/AP2σ^ang7^* (O-O’’)*, synj1^1/2^* (P-P’’), and *synj1^1/2^*; *wnd^2/3^* (Q-Q’’) immunolabeled for CSP (magenta) and HRP (green). The scale bar in Q’’ represents 10 µm. (**R**) Histogram showing bouton number from muscle 6/7 NMJ at A2 hemisegment for control (62.6 ± 2.88), *wnd^2/3^* (69 ± 3.25)*, AP2σ-^KG^/AP2σ^ang7^/AP2σ^ang7^* (129 ± 6.50)*, wnd^2/3^, AP2σ-^KG^/AP2σ^ang7^* (80.9 ± 5.04), *Elav-Gal4>UAS-AP2σ, AP2σ-^KG^/AP2σ^ang7^* (63.1 ± 4.92)*, synj1^1/2^* (154 ± 6.69), and *synj1^1/2^*; *wnd^2/3^* (92.2 ± 2.62). At least 8 NMJs from each genotype were used for quantification. (**S**) Histogram showing the average bouton area (µm^2^) from muscle 6/7 NMJ at A2 hemisegment for control (9.99 ± 0.55), *wnd^2/3^* (9.36 ± 0.45)*, AP2σ-^KG^/AP2σ^ang7^* (1.87 ± 0.19)*, wnd^2/3^, AP2σ-^KG^/AP2σ^ang7^* (7.77 ± 0.69), *Elav-Gal4>UAS-AP2σ, AP2σ-^KG^/AP2σ^ang7^* (7.94 ± 0.53)*, synj1^1/2^* (1.17 ± 0.09), and *synj1^1/2^*; *wnd^2/3^*(6.22 ± 0.15). At least 8 NMJs from each genotype were used for quantification. The error bars in R and S represent the standard error of the mean (SEM); the statistical analysis was done using one-way ANOVA followed by post hoc Tukey’s test. ***p<0.001; ns, not significant.

Elevated Wnd levels in the endocytic mutants suggested that synaptic phenotypes in these mutants are likely mediated by Wnd-dependent activation of JNK signaling. To directly test whether Wnd is required for the synaptic overgrowth phenotype, we generated double mutants of *wnd* and endocytic genes (*AP2* and *synj1*) and analyzed NMJ morphology. If loss of *wnd* suppresses the increased bouton number and satellite bouton formation observed in endocytic mutants, this would indicate that Wnd acts as an upstream activator of JNK signaling and is necessary for the synaptic overgrowth phenotype in endocytic mutants. Analysis of NMJ phenotypes in either *AP2* (*wnd^2^*^/3^, *AP2σ^KG/Ang7^*) or *synj1* (*synj1^1/2^; wnd^2/3^*) mutants revealed significant suppression of synaptic overgrowth, restoring bouton number, satellite boutons, and bouton area toward control values (Figure 4K-S and Figure 4 - figure supplement 2), indicating that Wnd upregulation is a major downstream effector of synaptic defects in endocytic mutants. These data indicate that an impaired endocytic pathway compromises Hiw function, leading to elevated Wnd and driving synaptic overgrowth.

### Endocytosis Restricts JNK Signaling in *Drosophila* Neurons

We next asked whether impaired Hiw function in endocytic mutants leads to activation of downstream Wnd/DLK signaling, a known regulator of synaptic growth through the JNK pathway in *Drosophila* (WAN *et al*. 2000; COLLINS *et al*. 2006). To directly monitor JNK signaling activation, we used the *puckered (puc)-lacZ* transcriptional reporter (Figure 5A) (MARTÍN-BLANCO *et al*. 1998; AGNES *et al*. 1999; ZHU *et al*. 2019). Both *AP2* mutants (Figure 5B-G) and *synj1* mutants showed significantly elevated *puc-lacZ* expression in the VNC (Figure 5E-G), indicating upregulation of JNK signaling in these endocytic mutants. This observation was further supported by a *TRE-DsRed* reporter, a synthetic construct that directly reports AP1/JNK activity (Figure 5A) (CHATTERJEE AND BOHMANN 2012). Consistent with a direct upregulation of JNK signaling, *TRE-DsRed* expression was robustly elevated in both *AP2* and *synj1* mutants (Figure 5H-M). Together, these results suggest that JNK signaling is upregulated in endocytic mutants and may contribute to synaptic overgrowth.

**Figure 5:**
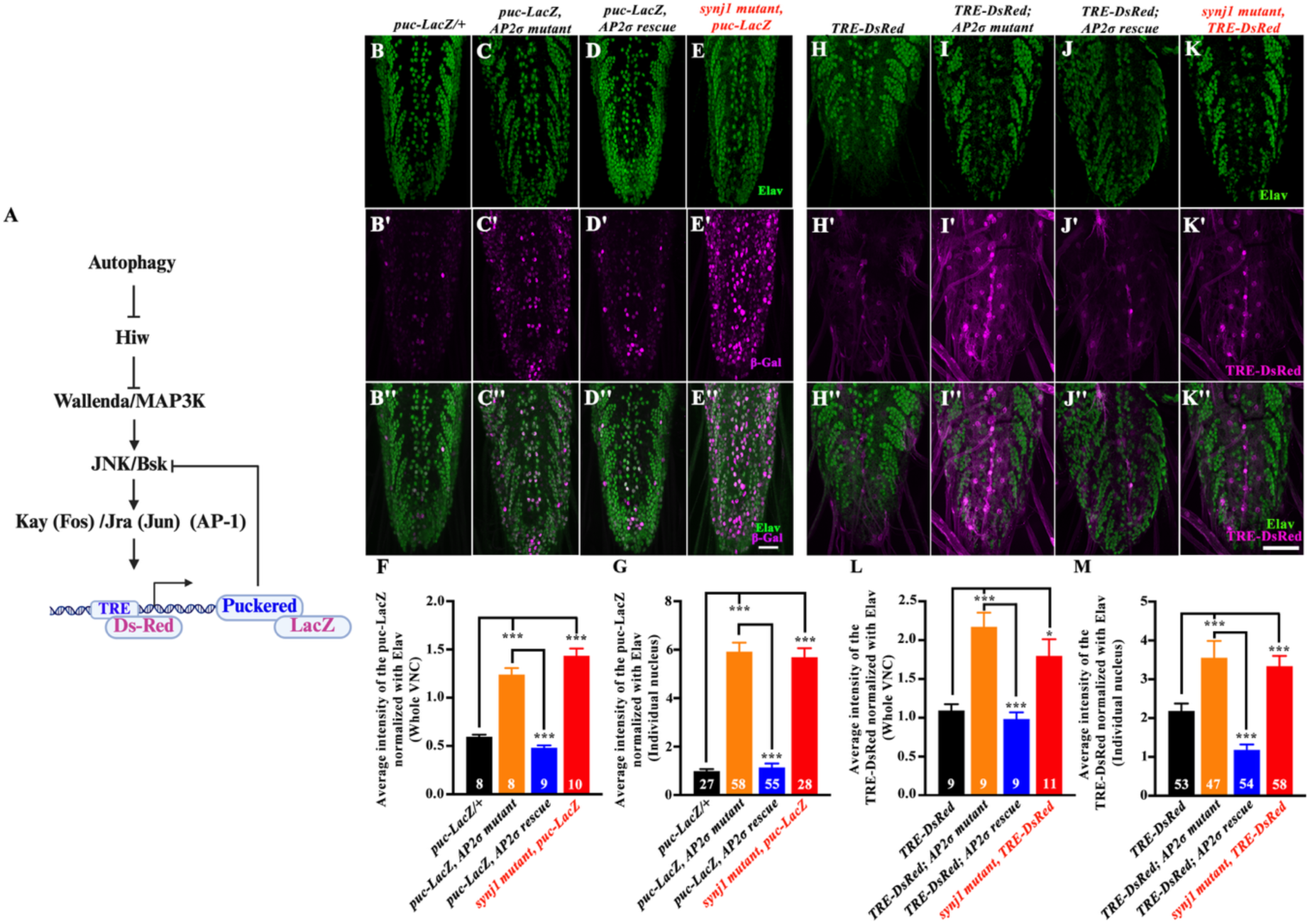
Endocytosis restricts hyperactivation of JNK signaling. (**A**) Schematic illustration of the JNK signaling pathway in *Drosophila.* Autophagy selectively degrades Hiw, leading to elevated levels of MAP3K Wallenda and eventually resulting in phosphorylation of Fos and Jun (AP-1). Phosphorylation of AP-1 results in activation of JNK signaling. TRE-DsRed (TRE is a synthetic promoter containing AP-1 binding sites) and puc-LacZ (puckered is a target gene of JNK signaling) were used as reporters to analyze the activation of the JNK signaling pathway. (**B-E’’**) Confocal images of third instar larval ventral nerve cord co-labelled for β-Gal (magenta) and Elav (green) in control (*puc-LacZ/+*) (B-B’’), *puc-LacZ, AP2σ-^KG^/AP2σ^ang7^*(C-C’’), *Elav-Gal4>UAS-AP2σ; puc-LacZ; AP2σ-^KG^/AP2σ^ang7^*, (D-D’’) and *synj1^1/2^; puc-LacZ/+* (E-E’’). The scale bar in E’’ represents 20 µm. (F) Histogram showing the β-Gal levels in the third instar larval ventral nerve cord in control (*puc-LacZ/+*) (0.59 ± 0.02), *puc-LacZ, AP2σ-^KG^/AP2σ^ang7^* (1.24 ± 0.06), *Elav-Gal4>UAS-AP2σ; puc-LacZ; AP2σ-^KG^/AP2σ^ang7^* (0.48 ± 0.02), and *synj1^1/2^; puc-LacZ/+* (1.43 ± 0.07). At least 8 VNCs of each genotype were used for the quantification. (G) Histogram showing the β-Gal levels in the individual neuronal nuclei in control (*puc-LacZ/+*) (0.99 ± 0.09), *puc-LacZ, AP2σ-^KG^/AP2σ^ang7^* (5.92 ± 0.37), *Elav-Gal4>UAS-AP2σ; puc-LacZ; AP2σ-^KG^/AP2σ^ang7^* (1.14 ± 0.16), and *synj1^1/2^; puc-LacZ/+* (5.69 ± 0.36). At least 27 nuclei of each genotype were used for the quantification. The error bars in F and G represent the standard error of the mean (SEM); the statistical analysis was done using one-way ANOVA followed by post hoc Tukey’s test. ***p<0.001; ns: not significant. (**H-K’’**) Confocal images of third instar larval ventral nerve cord co-labelled for TRE-DsRed (magenta) and Elav (green) in control (*TRE-DsRed/+*) (H-H’’), *TRE-DsRed/+; AP2σ-^KG^/AP2σ^ang7^* (I-I’’), *Elav-Gal4> UAS-AP2σ; TRE-DsRed/+; AP2σ-^KG^/AP2σ^ang7^* (J-J’’), and *TRE-DsRed/+, synj1^1/2^* (K-K’’). The scale bar in K’’ represents 20 µm. (**L**) Histogram showing the TRE-DsRed levels in the third instar larval ventral nerve cord in control (*TRE-DsRed/+*) (1.09 ± 0.08), *TRE-DsRed/+; AP2σ-^KG^/AP2σ^ang7^* (2.17 ± 0.18), *Elav-Gal4> UAS-AP2σ; TRE-DsRed/+; AP2σ-^KG^/AP2σ^ang7^* (0.98 ± 0.08), and *TRE-DsRed/+, synj1^1/2^* (1.80 ± 0.21). At least 9 VNCs of each genotype were used for quantification. (**M**) Histogram showing the TRE-DsRed levels in individual neuronal nuclei in control (*TRE-DsRed/+*) (2.19 ± 0.18), *TRE-DsRed/+; AP2σ-^KG^/AP2σ^ang7^* (3.56 ± 0.43), *Elav-Gal4> UAS-AP2σ; TRE-DsRed/+; AP2σ-^KG^/AP2σ^ang7^* (1.18 ± 0.13), and *TRE-DsRed, synj1^1/2^* (3.34 ± 0.26). At least 47 nuclei of each genotype were used for quantification. The error bars in L and M represent the standard error of the mean (SEM); the statistical analysis was performed using one-way ANOVA followed by Tukey’s post hoc test. ***p<0.001; *p<0.05; ns, not significant.

To directly test whether elevated JNK signaling is the primary pathway driving synaptic overgrowth in *AP2* mutant neurons, we genetically suppressed this pathway at two distinct levels: at the kinase level by expressing a dominant-negative form of the *Drosophila* JNK homolog Basket (*Bsk^DN^*), and at the transcriptional level by expressing a dominant-negative form of Kayak (*Kay^DN^* or *Fos^DN^*) (ERESH *et al*. 1997; WEBER *et al*. 2000; SANYAL *et al*. 2002; COLLINS *et al*. 2006; COLLINS AND DIANTONIO 2007b). We hypothesize that if JNK signaling were indeed the primary driver of synaptic overgrowth in *AP2* mutants, expression of either *Bsk^D^* or *Kay^DN^* should significantly suppress the synaptic defects characteristic of these mutants and phenocopy the suppression observed with loss-of-function mutations of *wnd*. A partial suppression by either construct would suggest that JNK contributes to, but does not fully account for, the overgrowth phenotype, consistent with the possibility of parallel pathways involved in regulation of synaptic growth such as the BMP signaling pathway (O’CONNOR-GILES *et al*. 2008; RODAL *et al*. 2011; CHOUDHURY *et al*. 2022). Conversely, if neither *Bsk^DN^* nor *Kay^DN^* affects bouton number or NMJ morphology in the *AP2* mutant background, this would indicate that JNK signaling is not required for overgrowth and that other pathways, such as BMP, are solely responsible. Analysis of synaptic phenotypes in these genetic manipulations (*Elav-Gal4>UAS-Bsk^DN^; AP2σ^KG/Ang7^* and *Elav-Gal4>UAS-Kay^DN^; AP2σ^KG/Ang7^*) revealed significant suppression of synaptic overgrowth in the endocytic mutants (Figure 6A-H, M-Q), supporting a causal role for JNK signaling in restricting synaptic growth. However, we found that neither *Bsk^DN^* nor *Kay^DN^* could completely suppress synaptic overgrowth (bouton number, satellite boutons, and bouton area) to the control levels, suggesting involvement of other parallel pathways, such as BMP signaling.

**Figure 6:**
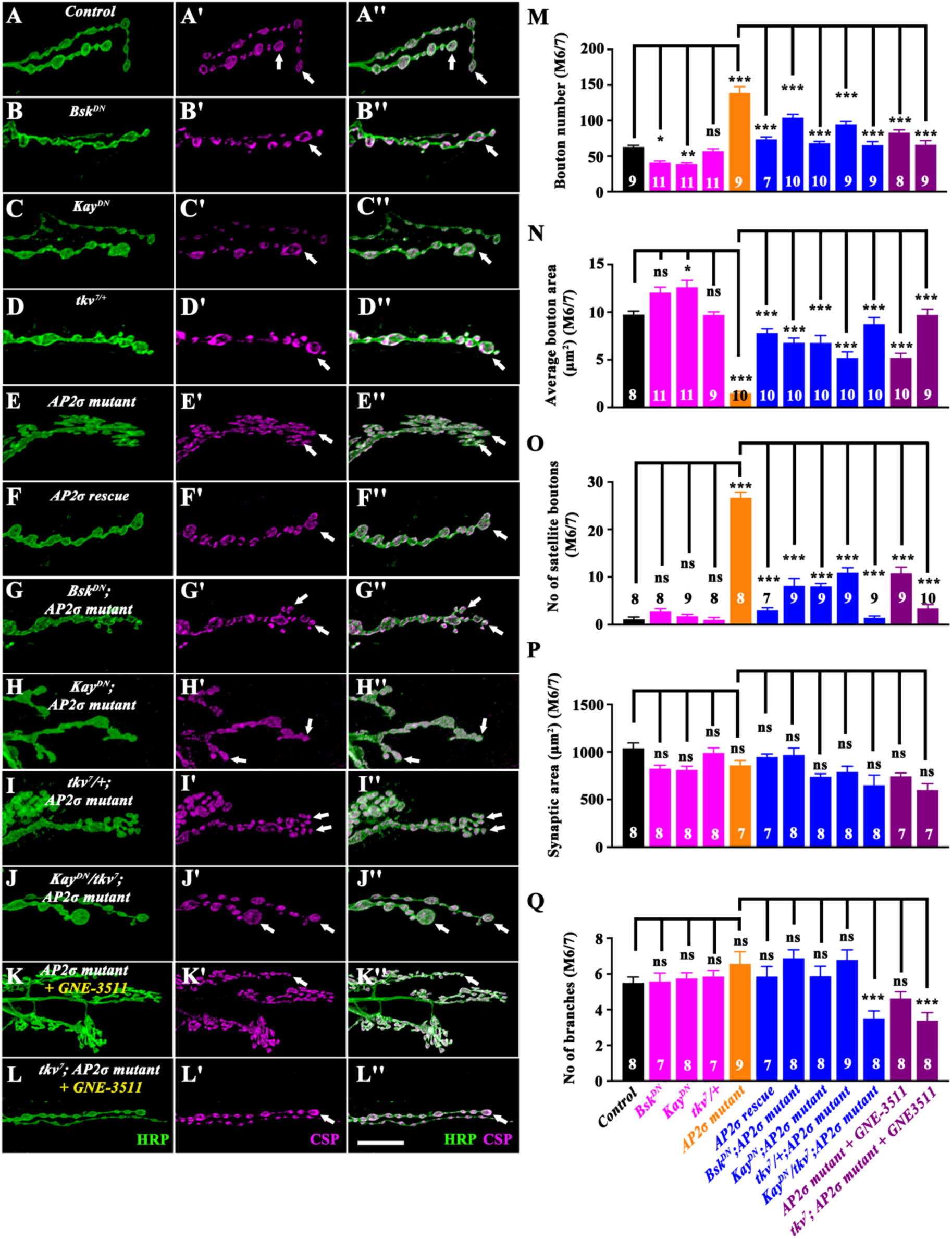
Elevated BMP and JNK signaling together promote synaptic overgrowth in endocytic mutants. (**A-L’’**) Confocal images of muscle 6/7 NMJ at A2 hemisegment showing synaptic growth in control (A-A’’), *Elav-Gal4>UAS-Bsk^DN^* (B-B’’), *Elav-Gal4>UAS-Kay^DN^* (C-C’’), *tkv^7/+^* (D-D’’)*, AP2σ-^KG^/AP2σ^ang7^* (E-E’’)*, Elav-Gal4> UAS-AP2σ, AP2σ-^KG^/AP2σ^ang7^* (F-F’’), *Elav-Gal4>UAS-Bsk^DN^; AP2σ-^KG^/AP2σ^ang7^* (G-G’’), *Elav-Gal4>UAS-Kay^DN^; AP2σ-^KG^/AP2σ^ang7^* (H-H’’), *tkv^7^/+; AP2σ-^KG^/AP2σ^ang7^*(I-I’’), *Elav-Gal4>UAS-Kay^DN^, tkv^7^; AP2σ-^KG^/AP2σ^ang7^*(J-J’’), *AP2σ-^KG^/AP2σ^ang7^* + DLK inhibitor (DLKi) (K-K’’), and *tkv^7^/+; AP2σ-^KG^/AP2σ^ang7^* + DLKi (L-L’’), immunolabeled for synaptic vesicle marker CSP (magenta) and neuronal membrane marker HRP (green). The scale bar in L’’ represents 10 µm. (M) Histogram showing bouton number in muscle 6/7 NMJ at A2 hemisegment in control (*w^1118^*) (62.9 ± 2.44), *Elav-Gal4>UAS-Bsk^DN^* (41.4 ± 2.19), *Elav-Gal4>UAS-Kay^DN^* (39 ± 1.95), *tkv^7^*^/+^ (57 ± 3.40), *AP2σ-^KG^/AP2σ^ang7^* (139 ± 8.80)*, Elav-Gal4>UAS-AP2σ, AP2σ-^KG^/AP2σ^ang7^* (73.6 ± 3.38), *Elav-Gal4>UAS-Bsk^DN^; AP2σ-^KG^/AP2σ^ang7^* (104 ± 4.59), *Elav-Gal4>UAS-Kay^DN^; AP2σ-^KG^/AP2σ^ang7^* (82.9 ± 3.03), *tkv^7^/+; AP2σ-^KG^/AP2σ^ang7^* (94.8 ± 3.91), *Elav-Gal4>UAS-Kay^DN^, tkv^7^; AP2σ-^KG^/AP2σ^ang7^* (65.6 ± 4.89), *AP2σ-^KG^/AP2σ^ang7^*+ DLKi (83.1 ± 3.94), and *tkv^7^/+; AP2σ-^KG^/AP2σ^ang7^* + DLKi (66.0 ± 5.87). At least 7 NMJs from each genotype were used for quantification. The error bar represents the standard error of the mean (SEM); the statistical analysis was done using one-way ANOVA followed by post hoc Tukey’s test. ***p<0.001; **p<0.01; *p<0.1; ns, not significant. (N) Histogram showing average bouton area (µm^2^) in muscle 6/7 NMJ at A2 hemisegment in control (*w^1118^*) (9.74 ± 0.35), *Elav-Gal4>UAS-Bsk^DN^* (12.1 ± 0.56), *Elav-Gal4>UAS-Kay^DN^* (12.6 ± 0.73), *tkv^7^*^/+^ (9.71 ± 0.32), *AP2σ-^KG^/AP2σ^ang7^* (1.51 ± 1.18)*, Elav-Gal4>UAS-AP2σ, AP2σ-^KG^/AP2σ^ang7^* (7.82 ± 0.4), *Elav-Gal4>UAS-Bsk^DN^; AP2σ-^KG^/AP2σ^ang7^*(6.81 ± 0.49), *Elav-Gal4>UAS-Kay^DN^; AP2σ-^KG^/AP2σ^ang7^* (6.79 ± 0.77), *tkv^7^/+; AP2σ-^KG^/AP2σ^ang7^* (5.19 ± 0.64), *Elav-Gal4>UAS-Kay^DN^, tkv^7^; AP2σ-^KG^/AP2σ^ang7^* (8.74 ± 0.69), *AP2σ-^KG^/AP2σ^ang7^*+ DLKi (5.20 ± 0.48), and *tkv^7^/+; AP2σ-^KG^/AP2σ^ang7^* + DLKi (9.71 ± 0.59). At least 8 NMJs from each genotype were used for quantification. The error bar represents the standard error of the mean (SEM); the statistical analysis was done using one-way ANOVA followed by post hoc Tukey’s test. ***p<0.001; *p<0.05; ns, not significant. **(O)** Histogram showing number of satellite boutons in muscle 6/7 NMJ at A2 hemisegment in control (*w^1118^*) (1.13 ± 0.48), *Elav-Gal4>UAS-Bsk^DN^* (2.75 ± 0.59), *Elav-Gal4>UAS-Kay^DN^* (1.78 ± 0.4), *tkv^7^*^/+^ (1 ± 0.5), *AP2σ-^KG^/AP2σ^ang7^* (26.6 ± 1.16)*, Elav-Gal4>UAS-AP2σ, AP2σ-^KG^/AP2σ^ang7^* (3.00 ± 0.53), *Elav-Gal4>UAS-Bsk^DN^; AP2σ-^KG^/AP2σ^ang7^* (8.11 ± 1.59), *Elav-Gal4>UAS-Kay^DN^; AP2σ-^KG^/AP2σ^ang7^* (8.00 ± 0.60), *tkv^7^/+; AP2σ-^KG^/AP2σ^ang7^* (10.9 ± 1.06), *Elav-Gal4>UAS-Kay^DN^, tkv^7/+^; AP2σ-^KG^/AP2σ^ang7^* (1.44 ± 0.37), *AP2σ-^KG^/AP2σ^ang7^* + DLKi (10.8 ± 1.29), and *tkv^7^/+; AP2σ-^KG^/AP2σ^ang7^* + DLKi (3.40 ± 0.80). At least 7 NMJs from each genotype were used for quantification. The error bar represents the standard error of the mean (SEM); the statistical analysis was done using one-way ANOVA followed by post hoc Tukey’s test. ***p<0.001; ns, not significant. **(P)** Histogram showing synaptic area (µm^2^) in muscle 6/7 NMJ at A2 hemisegment in control (*w^1118^*) ( 1038 ± 57.7), *Elav-Gal4>UAS-Bsk^DN^* (825 ± 34.4), *Elav-Gal4>UAS-Kay^DN^* (811 ± 37.7), *tkv^7^*^/+^ (989 ± 54.5), *AP2σ-^KG^/AP2σ^ang7^* (858 ± 52.3)*, Elav-Gal4>UAS-AP2σ, AP2σ-^KG^/AP2σ^ang7^* (947 ± 31.1), *Elav-Gal4>UAS-Bsk^DN^; AP2σ-^KG^/AP2σ^ang7^* (969 ± 72.2), *Elav-Gal4>UAS-Kay^DN^; AP2σ-^KG^/AP2σ^ang7^* (740 ± 31.8), *tkv^7^/+; AP2σ-^KG^/AP2σ^ang7^* (790 ± 58.8), *Elav-Gal4>UAS-Kay^DN^, tkv^7^; AP2σ-^KG^/AP2σ^ang7^* (651 ± 106), *AP2σ-^KG^/AP2σ^ang7^* + DLKi (744 ± 34.3), and *tkv^7^/+; AP2σ-^KG^/AP2σ^ang7^* + DLKi (600 ± 67.8). At least 7 NMJs from each genotype were used for quantification. The error bar represents the standard error of the mean (SEM); the statistical analysis was done using one-way ANOVA followed by post hoc Tukey’s test. ns, not significant. **(Q)** Histogram showing the number of branches in muscle 6/7 NMJ at A2 hemisegment in in control (*w^1118^*) (5.50 ± 0.32), *Elav-Gal4>UAS-Bsk^DN^* (5.57 ± 0.48), *Elav-Gal4>UAS-Kay^DN^* (5.75 ± 0.31), *tkv^7^*^/+^ (5.86 ± 0.34), *AP2σ-^KG^/AP2σ^ang7^* (6.56 ± 0.68)*, Elav-Gal4>UAS-AP2σ, AP2σ-^KG^/AP2σ^ang7^* (5.86 ± 0.55), *Elav-Gal4>UAS-Bsk^DN^; AP2σ-^KG^/AP2σ^ang7^* (6.88 ± 0.48), *Elav-Gal4>UAS-Kay^DN^; AP2σ-^KG^/AP2σ^ang7^* (5.88 ± 0.54), *tkv^7^/+; AP2σ-^KG^/AP2σ^ang7^* (6.78 ± 0.57), *Elav-Gal4>UAS-Kay^DN^, tkv^7^; AP2σ-^KG^/AP2σ^ang7^* (3.50 ± 0.42), *AP2σ-^KG^/AP2σ^ang7^* + DLKi (4.63 ± 0.37), and *tkv^7^/+; AP2σ-^KG^/AP2σ^ang7^* + DLKi (3.38 ± 0.46). At least 7 NMJs from each genotype were used for quantification. The error bar represents the standard error of the mean (SEM); the statistical analysis was done using one-way ANOVA followed by post hoc Tukey’s test. ***p<0.001; ns, not significant.

Endocytosis is also known to attenuate BMP signaling and restrict synaptic growth (WU *et al*. 2005b; O’CONNOR-GILES *et al*. 2008; GLEASON *et al*. 2014; JACOMIN *et al*. 2015; EHRLICH 2016; GLEASON *et al*. 2017). Like the JNK pathway, BMP pathway suppression (*tkv^7^*^/+^ or *wit^A12^*^/+^) alone is not sufficient to rescue the synaptic overgrowth phenotypes in endocytic mutants to the control level (Figure 6I-I’’ and M-Q) (CHOUDHURY *et al*. 2022). Consistent with these data, homozygous mutants of the BMP pathway receptor *wit* (*wit^B11/A12^, AP2σ^KG/Ang7^*) also did not completely suppress the clustering and synaptic overgrowth phenotypes in endocytic mutants (Figure 6 - figure supplement 1), suggesting involvement of the JNK pathway in suppressing synaptic growth.

We next asked whether JNK and BMP pathways function in parallel to restrict synapse growth. We found that animals expressing GFP-Hiw in the endocytic mutants did not suppress the elevated p-MAD levels in *AP2* mutants (Figure 6 - figure supplement 2), suggesting that JNK signaling operates independently of the BMP pathway. Next, we co-expressed a dominant-negative form of *Kay* alongside a loss-of-function allele of *tkv* (*tkv^7/+^*), a BMP receptor, in *AP2* mutants. Strikingly, this combination fully suppressed bouton number, area, and satellite boutons to wild-type levels (Figure 6J-J’’ and M-Q).

To strengthen our observations that elevated JNK signaling functionally contributes to synaptic overgrowth in endocytic mutants, we treated larvae with GNE-3511 (DLKi), a pharmacological inhibitor of Wnd/DLK-1 (Figure 6 - figure supplement 3) (PATEL *et al*. 2015; HAYNE AND DIANTONIO 2022). Consistent with the genetic data, DLKi treatment did not fully suppress the synaptic overgrowth phenotypes to control levels. However, when *tkv^7^*^/+^; *AP2σ^KG/Ang7^* larvae were grown in DLKi-containing media, we found a complete suppression of synaptic overgrowth phenotypes (Figure 6K-L’’ and M-Q). Together, these findings strongly support the role of JNK signaling in mediating synaptic growth and indicate that both JNK and BMP signaling pathways function in parallel to regulate synaptic growth.

### Loss of Rab11 Promotes Formation of Hiw/Phr1 Assemblies and Activates JNK Pathway Akin to Endocytic Mutants

While Hiw accumulates in endocytic mutants, autophagy-deficient mutants show no such cytoplasmic Hiw assemblies despite elevated total Hiw levels (Figure 2 - figure supplement 2). This observation suggests that reduced Hiw degradation alone is insufficient for puncta formation and that trafficking defects could be a critical determinant of Hiw accumulation in endocytic mutants. Hence, to directly test whether Hiw trafficking involves the endosomal system, we neuronally expressed dominant-negative (DN) forms of the small GTPases Rab5, Rab7, or Rab11, which regulate different steps of endocytosis and recycling. While expression of DN-Rab5 (Rab5^S43N^) or DN-Rab7 (Rab7^T22N^) did not affect Hiw localization in neurons, expression of DN-Rab11 (Rab11^S25N^) resulted in large assemblies and accumulation of Hiw within neuronal cell bodies, phenocopying the endocytic mutants (Figure 7A-E).

**Figure 7:**
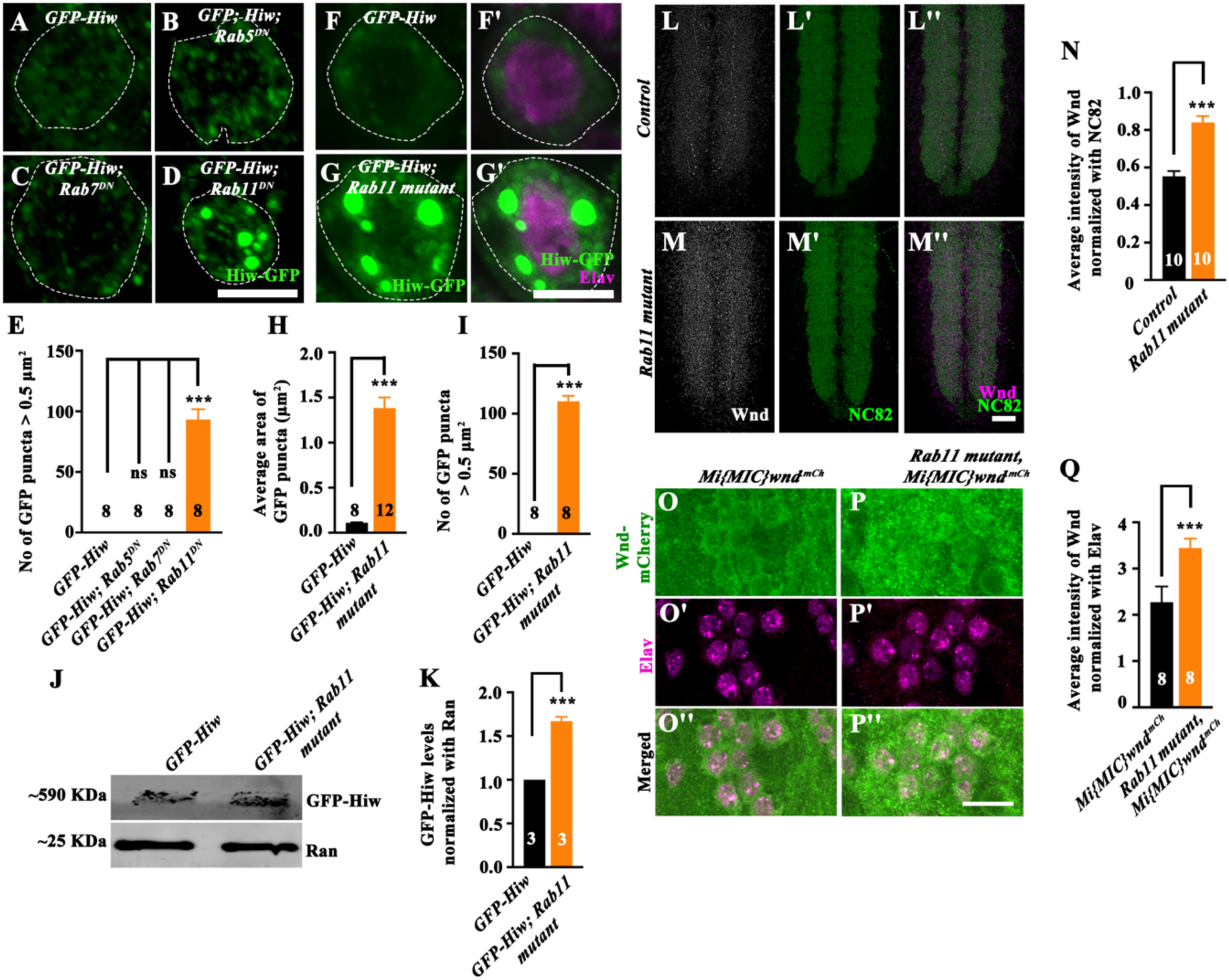
Loss of Rab11 induces Highwire accumulation in neuronal cell body. (**A-D**) Confocal images of the neuronal cell body labelled for GFP-Hiw (green) in control (*Elav-Gal4>UAS-GFP-Hiw*) (A), *Elav-Gal4>UAS-GFP-Hiw; UAS-Rab5^DN^* (B), *Elav-Gal4>UAS-GFP-Hiw; UAS-Rab7^DN^* (C), and *Elav-Gal4>UAS-GFP-Hiw; UAS-Rab11^DN^* (D). The scale bar in D represents 5 µm. (**E**) Histogram showing the number of GFP-Hiw puncta with area greater than 0.5 μm^2^ in control (*Elav-Gal4>UAS-GFP-Hiw*) (0.00 ± 0.00), *Elav-Gal4>UAS-GFP-Hiw; UAS-Rab5^DN^* (0.00 ± 0.00), *Elav-Gal4>UAS-GFP-Hiw; UAS-Rab7^DN^* (0.00 ± 0.00), *Elav-Gal4>UAS-GFP-Hiw; UAS-Rab11^DN^* (93.3 ± 8.52). At least 8 VNCs of each genotype were used for the quantification. The error bar represents the standard error of the mean (SEM); the statistical analysis was done using one-way ANOVA followed by post hoc Tukey’s test. ***p<0.001, ns: not significant (**F-G’**) Confocal images of the neuronal cell body co-labeled for GFP-Hiw (green) and Elav (magenta) in control (*Elav-Gal4>UAS-GFP-Hiw*) (F-F’), and *Elav-Gal4>UAS-GFP-Hiw; Rab11^ex2/93Bi^* (G-G’). The scale bar in G’ represents 5 µm. (H) Histogram showing the average area (µm^2^) of the GFP-Hiw puncta in the neuronal cell body of control (*Elav-Gal4>UAS-GFP-Hiw*) (0.11 ± 0.01) and *Elav-Gal4>UAS-GFP-Hiw; Rab11^ex2/93Bi^* (1.38 ± 0.11). At least 8 VNCs of each genotype were used for quantification. The error bar represents the standard error of the mean (SEM); the statistical analysis was done using Student’s t-test. ***p<0.001. (I) Histogram showing the number of GFP-Hiw puncta with area greater than 0.5 μm^2^ in the neuronal cell body of control (*Elav-Gal4>UAS-GFP-Hiw*) (0.00 ± 0.00) and *Elav-Gal4>UAS-GFP-Hiw; Rab11^ex2/93Bi^* (110 ± 4.59). At least 8 VNCs of each genotype were used for the quantification. The error bar represents the standard error of the mean (SEM); the statistical analysis was done using Student’s t-test. ***p<0.001. (J) Western blot showing the level of GFP-Hiw in control (*Elav-Gal4>UAS-GFP-Hiw*) and *Elav-Gal4>UAS-GFP-Hiw; Rab11^ex2/93Bi^*. Ran was used as the loading control. (K) Histogram showing the level of GFP-Hiw in control (*Elav-Gal4>UAS-GFP-Hiw*) (1.00 ± 0.00) and *Elav-Gal4>UAS-GFP-Hiw; Rab11^ex2/93Bi^* (1.67 ± 0.05). Three independent Western blots were used for quantification. The error bar represents the standard error of the mean (SEM); the statistical analysis was done using Student’s t-test. ***p<0.001. (**L-M’’**) Confocal images of third-instar larval ventral nerve cord co-labeled for Wnd (magenta) and NC82 (green) in control (L-L’’), and *Rab11^ex2/93Bi^* (M-M’’). The scale bar in M’’ represents 10 µm. (**N**) Histogram showing the average fluorescence intensity of Wnd in control (0.55 ± 0.02), and *Rab11^ex2/93Bi^* (0.8 ± 0.03). At least 10 VNCs of each genotype were used for the quantification. The error bar represents the standard error of the mean (SEM); the statistical analysis was done using one-way ANOVA followed by post hoc Tukey’s test. ***p<0.001. (**O-P’’**) Confocal images of third-instar larval ventral nerve cord co-labeled for Wnd (green) and Elav (magenta) in control (*Mi{MIC}wnd^.mCh^)* (O-O’’), and *Rab11^ex2/93Bi^; Mi{MIC}wnd^.mCh^* (P-P’’). The scale bar in P’’ represents 10 µm. (**Q**) Histogram showing the average fluorescence intensity of Wnd in control (*Mi{MIC}wnd^.mCh^*) (2.27 ± 0.33), and *Rab11^ex2/93Bi^*; *Mi{MIC}wnd^.mCh^* ( 3.67 ± 0.2). At least 8 VNCs of each genotype were used for quantification. The error bar represents the standard error of the mean (SEM); the statistical analysis was done using one-way ANOVA followed by post hoc Tukey’s test. ***p<0.001; ns, not significant.

Consistent with these findings, *Rab11* mutants also displayed both Hiw protein assemblies and elevated levels of Hiw substrate Wnd/DLK (Figure 7F-Q and Figure 7 - figure supplement 1). Strikingly, synaptic overgrowth in *Rab11* mutants was significantly suppressed by neuronal expression of dominant-negative *Bsk^DN^* or dominant-negative *Kay^DN^*(Figure 8A-I), suggesting that JNK pathway activation lies downstream of Rab11 activity. These genetic epistatic interactions suggest that the disrupted Rab11-dependent recycling impedes proper localization of Hiw, thereby triggering aberrant JNK signaling and synaptic overgrowth. Given that Rab11-positive recycling endosomes are reduced in both *AP2* and *synj1* mutants (CHOUDHURY *et al*. 2022), these findings suggest that Rab11-dependent endosomal recycling is required for the localization/trafficking of Hiw, and its dysfunction contributes to Hiw accumulation in cell bodies, functional inactivation, and subsequent JNK pathway hyperactivation in endocytic mutants and synaptic overgrowth.

**Figure 8:**
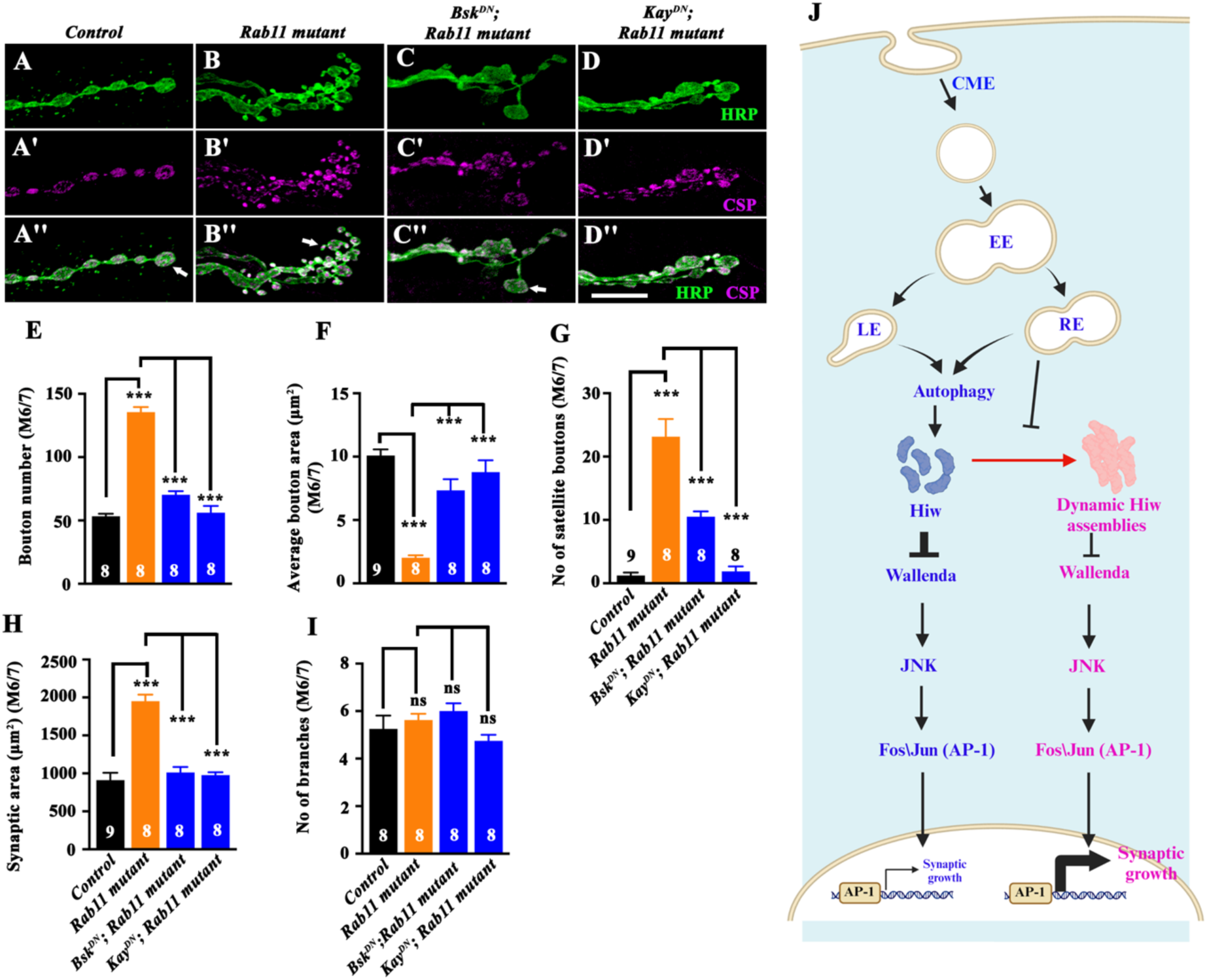
Endocytosis restricts JNK and BMP signaling through Rab11 in a parallel pathway. (**A-D’’**) Confocal images of muscle 6/7 NMJ showing the synaptic growth in control (A-A’’), *Rab11^ex2/93Bi^* (B-B’’)*, Elav-Gal4>UAS-Bsk^DN^; Rab11^ex2/93Bi^* (C-C’’), and *Elav-Gal4>UAS-Kay^DN^; Rab11^ex2/93Bi^* (D-D’’), immunolabeled for CSP (magenta) and HRP (green). The scale bar in D’’ represents 10 µm. (E) Histogram showing bouton number from muscle 6/7 NMJ at A2 hemisegment in control (53.3 ± 2.03), *Rab11^ex2/93Bi^* (136 ± 4.02)*, Elav-Gal4>UAS-Bsk^DN^; Rab11^ex2/93Bi^* (70.3 ± 2.91) and *Elav-Gal4>UAS-Kay^DN^; Rab11^ex2/93Bi^* (56.1 ± 5.32). At least 8 NMJs from each genotype were used for quantification. The error bar represents the standard error of the mean (SEM); the statistical analysis was done using one-way ANOVA followed by post hoc Tukey’s test. ***p<0.001. (F) Histogram showing the average bouton area (µm^2^) from muscle 6/7 NMJ at A2 hemisegment in control (10.1 ± 0.48), *Rab11^ex2/93Bi^* (2.02 ± 0.19)*, Elav-Gal4>UAS-Bsk^DN^; Rab11^ex2/93Bi^* (7.33 ± 0.88), and *Elav-Gal4>UAS-Kay^DN^; Rab11^ex2/93Bi^* (8.78 ± 0.93). At least 8 NMJs from each genotype were used for quantification. The error bar represents the standard error of the mean (SEM); the statistical analysis was done using one-way ANOVA followed by post hoc Tukey’s test. ***p<0.001. (G) Histogram showing the number of satellite boutons in muscle 6/7 NMJ at A2 hemisegment in control (1.22 ± 0.46), *Rab11^ex2/93Bi^* (23.1 ± 2.79)*, Elav-Gal4>UAS-Bsk^DN^; Rab11^ex2/93Bi^* (10.5 ± 0.84), and *Elav-Gal4>UAS-Kay^DN^; Rab11^ex2/93Bi^* (1.88 ± 0.79). At least 8 NMJs from each genotype were used for quantification. The error bar represents the standard error of the mean (SEM); the statistical analysis was done using one-way ANOVA followed by post hoc Tukey’s test. ***p<0.001. (H) Histogram showing the synaptic area (µm^2^) in muscle 6/7 NMJ at A2 hemisegment in the control (910 ± 9.69) and *Rab11^ex2/93Bi^* (1951 ± 86.2), *Elav-Gal4>UAS-Bsk^DN^; Rab11^ex2/93Bi^* (1011 ± 72.8), *Elav-Gal4>UAS-Kay^DN^; Rab11^ex2/93Bi^* (978 ± 36.8). At least 8 NMJs were used for quantification. The error bar represents the standard error of the mean (SEM), and statistical analysis was done using Student’s t-test. ***p<0.001. (I) Histogram showing the number of branches in muscle 6/7 NMJ at A2 hemisegment in the control (5.25 ± 0.56) and *Rab11^ex2/93Bi^* (5.63 ± 0.26), *Elav-Gal4>UAS-bsk^DN^; Rab11^ex2/93Bi^* (6 ± 0.33), *Elav-Gal4>UAS-kay^DN^; Rab11^ex2/93Bi^* (4.75 ± 0.25). At least 8 NMJs were used for quantification. The error bar represents the standard error of the mean (SEM), and statistical analysis was done using Student’s t-test. ns: not significant. (J) Proposed model of synaptic growth signaling regulation. Endocytosis regulates Rab11-positive recycling endosomes, which in turn modulate three partially independent pathways to attenuate synaptic growth. First, Rab11 regulates endosomal trafficking of Hiw; when this trafficking is impaired, Hiw undergoes accumulation into large, nonfunctional assemblies, thereby hyperactivating JNK signaling. Second, reduced Rab11 impairs autophagosome formation and autophagy, leading to elevated Hiw levels that accumulate as dynamic and reversible puncta in the endocytic mutants. Third, Rab11 regulates BMP receptor trafficking, and its reduction increases pMad levels. These pathways suggest that endocytosis restricts synaptic growth by constraining BMP and JNK signaling through a parallel Rab11-dependent endosomal trafficking pathway, with Hiw protein assembly as a key regulatory mechanism.

## Discussion

Endocytosis is increasingly recognized as a key modulator of neuronal signaling beyond its canonical roles in vesicle recycling (COSKER AND SEGAL 2014). Here, we identify a previously unrecognized function of endocytic trafficking in constraining synaptic growth by preventing aberrant accumulation of E3 ubiquitin ligase Highwire. When endocytic recycling is compromised, Hiw trafficking is disrupted, leading to its accumulation and functional inactivation. This loss in Hiw activity causes Wnd/DLK upregulation and JNK hyperactivation, resulting in synaptic overgrowth independent of autophagic dysfunction. Moreover, we identify Rab11 as a critical regulator of Hiw localization and trafficking.

### A Role for Endocytosis in Regulating Synaptic Stability by Modulating Hiw/Phr1 Function

Prior studies proposed that synaptic overgrowth observed in autophagy mutants arises from reduced Hiw turnover (SHEN AND GANETZKY 2009). However, paradoxically, *AP2* and *synj1* mutants exhibit Hiw accumulation as puncta in the neuronal cell bodies, aberrantly elevated JNK signaling, and synaptic overgrowth, phenotypes that are consistent with reduced Hiw/Phr1 function.

Our data reveal that endocytic proteins serve a broader role beyond synaptic vesicle recycling. Endocytosis likely regulates the localization/trafficking of Hiw/Phr1. We provide several lines of evidence to support this conclusion: a) acute disruption of CME via temperature-sensitive paralytic mutant *shibire^ts^* causes Hiw mislocalization and puncta formation; b) loss of recycling endosomal GTPase Rab11 phenocopies these defects, directly implicating endosomal recycling in Hiw regulation; and c) the puncta represent non-vesicular (Rab5-, Rab7-, and Rab11-positive) dynamic protein accumulation, rather than degradation intermediates, indicating that Hiw is sequestered rather than degraded under endocytic/trafficking block. Thus, endocytosis mediates Hiw trafficking to its functional sites, such as synapses (XIONG *et al*. 2010; XIONG *et al*. 2012b; GRILL *et al*. 2016) (Figure 1 – figure supplement 1E-I and Figure 7 - figure supplement 1), and protects it from aberrant sequestration, a mechanism consistent with broader observations that disruption of membrane trafficking can mislocalize critical signaling molecules in neurons (LEITCH *et al*. 2014). Since neuronal morphology and function depend on spatially restricted signaling, CME likely contributes to synaptic stability by protecting key regulatory complexes, such as Hiw/Phr1, from spatial miscompartmentalization.

These findings place Hiw/Phr1 within a growing class of neuronal proteins that undergo dynamic biomolecular assemblies to regulate synaptic structure and function. For example, postsynaptic scaffolds such as PSD-95 and SynGAP form LLPS condensates to assemble postsynaptic densities and tune synaptic plasticity (ZENG *et al*. 2016). Similarly, presynaptic scaffolds such as SYD-2/Liprin-α and ELKS undergo LLPS to establish active zones in *C. elegans* and mammals (MCDONALD *et al*. 2020). While our data do not conclusively establish Hiw assemblies as bona fide condensates, our observation that Hiw puncta are dynamic assemblies highlights an underappreciated layer of regulation in which membrane trafficking intersects with mechanisms that spatially gate enzymatic activity, thereby regulating neuronal signaling.

### Autophagy-Endocytosis Crosstalk Regulates Highwire Compartmentalization and Synaptic Homeostasis

While autophagy facilitates the turnover of synaptic signaling proteins (SHEN AND GANETZKY 2009; HOFFMANN-CONAWAY *et al*. 2020), endocytosis controls autophagic flux by regulating membrane supply and the trafficking of autophagy-related proteins (BIRGISDOTTIR AND JOHANSEN 2020; ANDRES-ALONSO *et al*. 2021). Endosomes are a primary membrane source for autophagosome formation (RAVIKUMAR *et al*. 2010; PURI *et al*. 2013; BIRGISDOTTIR AND JOHANSEN 2020), linking autophagic processes to endocytic flux. Endocytic proteins such as AP2 and Synj1 also modulate autophagy through endocytosis-independent mechanisms, particularly at the synaptic terminals (VANHAUWAERT *et al*. 2017; BERA *et al*. 2020). Thus, autophagy defects in endocytic mutants may arise from defective membrane supply and direct regulatory functions of endocytic proteins on autophagic machinery. Our analysis reveals that endocytosis and endosomal trafficking, but not autophagy, are essential to prevent aberrant accumulation of Hiw into large puncta within neuronal cell bodies. These puncta, solubilized by 1,6-HD (KROSCHWALD *et al*. 2017) (Figure 2E-Q), are not part of the endosomes or stress granules and are absent in autophagy-deficient animals (Figure 2 - figure supplement 1 and Figure 2 - figure supplement 2). In autophagy mutants such as *Atg1^Δ3D^*, we observed elevated steady-state Hiw levels, consistent with impaired degradation, but no Hiw puncta formation. Similarly, pharmacological inhibition of autophagy increased total Hiw abundance but did not promote Hiw assemblies. Thus, autophagy determines Hiw abundance, while endocytic trafficking determines its spatial distribution and compartmentalization.

Our data showing that acute disruption of CME in *shi^ts^*mutants was sufficient to trigger rapid Hiw assembly into large cytoplasmic puncta (Figure 3A-N) further support the view that endocytosis maintains Hiw in a soluble, spatially accessible pool. Autophagy alone is insufficient to control Hiw/Phr1 distribution. This separation of roles supports a two-tiered control of Hiw/Phr1 regulation: autophagy limits Hiw/Phr1 levels, while endocytosis prevents aberrant assembly of Hiw/Phr1. While we do not rule out indirect contributions of autophagy impairment under these conditions, the absence of Hiw puncta in autophagy-deficient animals strongly argues that autophagic inhibition alone is insufficient to induce Hiw assemblies. These results reveal that endocytosis maintains synaptic homeostasis by regulating Hiw in a spatially accessible manner by maintaining a soluble Hiw pool. While autophagy controls total Hiw protein levels, endocytic trafficking prevents its aberrant sequestration into dynamic assemblies, thereby restricting Hiw-mediated synaptic growth.

### Rab11-Dependent Endosomal Recycling: Essential Spatial Control Node for Synaptic Growth Regulation

Rab11-positive recycling endosomes serve as dynamic platforms that integrate endocytosis, membrane recycling, and autophagy. They not only contribute to autophagosome biogenesis by supplying membrane and trafficking autophagy-related proteins but also play essential roles in receptor trafficking and synaptic homeostasis (ULLRICH *et al*. 1996; GREEN *et al*. 1997; WILCKE *et al*. 2000; BARTZ *et al*. 2003; LONGATTI *et al*. 2012; LONGATTI AND TOOZE 2012; ZULKEFLI *et al*. 2019). In *Drosophila*, Rab11 cooperates with Nwk to regulate BMP receptor recycling, actin dynamics, and synaptic growth signaling at the NMJ (RODAL *et al*. 2008). Similarly, *AP2* and *Synj1* mutants genetically interact with *Rab11* mutants, and Rab11 levels are significantly reduced at the *AP2* mutant synapses, implicating endocytic recycling as a key signaling integration node between CME and autophagic flux (CHOUDHURY *et al*. 2022).

In *C. elegans*, the E3 ubiquitin ligase RPM-1 regulates synaptic development through interactions with the Rab GTPase GLO-1 and its GEF GLO-4 via late endosomes, suggesting that endosomal pathways help spatially restrict RPM-1 function within neurons (GRILL *et al*. 2007; OPPERMAN AND GRILL 2014b). While the specific endosomal compartment may differ across species, our findings in *Drosophila* similarly implicate endosomal trafficking in the localization and function of Hiw, the RPM-1 ortholog. Our data reveal that Rab11-mediated endosomal recycling is an essential trafficking event preventing Hiw accumulation.

A recent finding suggests that Rab11 is a critical mediator in the anterograde transport and sorting of Wnd to axon terminals for degradation, and that disruptions of this process lead to Wnd accumulation, excessive stress signaling, and neuronal degeneration (KIM *et al*. 2024). Our study offers an alternative or additional mechanism for these observations and supports the idea that loss of Rab11 renders Hiw inactive by promoting accumulation, thereby leading to elevated Wnd levels. Together with CME, Rab11-positive endosomes form a trafficking axis that regulates the spatial dynamics and functional availability of Hiw at synapses, preventing overactivation of JNK activity and restricting synapse overgrowth. JNK signaling has long been implicated in regulating synaptic expansion, plasticity, and dendritic architecture across species, from *Drosophila* to mammals (WU *et al*. 2005a; ABRAMS *et al*. 2008; OPPERMAN AND GRILL 2014a). The genetic suppression of JNK signaling by blocking the components of the JNK pathway or Wnd mutants demonstrates that synaptic overgrowth in endocytic mutants is directly mediated by JNK pathway activation downstream of Wnd (Figures 4K-S and 6). Notably, combined inhibition of JNK and BMP pathways in endocytic or Rab11 mutants fully restores synaptic morphology, indicating that JNK acts in parallel, not simply downstream, of BMP in restricting synaptic growth, supporting emerging models that growth-promoting pathways integrate within a context-dependent, multi-pathway framework. We suggest a dual safeguarding model (Figure 8J) in which endocytosis and Rab11-mediated recycling endosomes spatially govern Hiw distribution, preventing its sequestration into larger, dynamic protein assemblies. Autophagy provides complementary support for Hiw turnover. Together, these mechanisms maintain a critical threshold for Wnd-JNK signaling in neurons and restrict synaptic overgrowth. Although GFP-Hiw forms higher-order, reversible assemblies under conditions of impaired endocytosis or vesicular trafficking, further investigation is required to establish whether these structures represent bona fide biomolecular condensates. Nevertheless, our findings highlight a potential link between endocytic trafficking and the regulation of Hiw-JNK signaling that extends beyond developmentally regulated synaptic growth. This relationship may also have implications for axon regeneration and neuronal maintenance, as defects in endocytosis and endosomal trafficking have been associated with enhanced JNK signaling, neuronal loss, and synaptic degeneration.

## Materials and Methods

### Fly Stocks

All the flies were grown and maintained at 25^°^C in a standard cornmeal medium as described previously (CHOUDHURY *et al*. 2016). *w^1118^* was used as a control unless otherwise mentioned. Both males and females were used for experiments. The *AP2σ* mutant alleles used were *AP2σ^ang7^*, a hypomorphic allele generated through P-element mobilization of *Synd^EP877^*located 2416 bp upstream of the Syndapin ORF, and *AP2σ^KG0245^*, a null allele of AP2σ which has a P-element insertion in the third exon of the AP2σ ORF (CHOUDHURY *et al*. 2016). The following fly stocks were obtained from BDSC: *AP2σ^KG02457^* (BL13478), *UAS-GFP-mCherry-Atg8a* (BL37749), *UAS-GFP-Atg8a* (BL52005), *UAS-GFP-Hiw* (BL51639), *UAS-AP2β RNAi* (BL28328), *UAS-bsk^DN^* (BL6409), *UAS-kay^DN^* (BL7214*), synj1^1^* (BL24883)*, synj1^2^* (BL24884), *tkv^7^*(BL3242), *wnd^2^*, *wnd^3^* (BL51999) (COLLINS *et al*. 2006), TRE-DsRed (BL59011), *UAS-Rab5^S43N^* (BL42703), *UAS-Rab7^T22N-YFP^* (BL9778), *UAS-Rab11^S25N-YFP^* (BL23261), *witA^12^*(BL5173), *wit^B11^*(BL5174), *hiw^ΔN^* (BL51637), and *Mi{MIC}Wnd^.mCh^* (BL39657). Other lines used in the study are: *atg1^Δ3D^ (SCOTT et al. 2004)*, *UAS-AP2α RNAi* (GD15566), *Rab11^93Bi^*, *Rab11^ex2^*(KHODOSH *et al*. 2006), and *puc-lacZ* (RING AND MARTINEZ ARIAS 1993).

The *shi^ts1^* and *shi^ts2^* temperature-sensitive mutant fly stocks were maintained at 18°C, and experimental crosses were set up at the permissive temperature of 22°C. Once the progeny reached the wandering third instar larval stage, larvae were shifted to the non-permissive temperature of 29°C for 3 hours and 6 hours to induce acute disruption of dynamin-mediated endocytosis (KOSAKA AND IKEDA 1983; DICKMAN *et al*. 2006; KROLL *et al*. 2015).

The DLK inhibitor GNE-3511 (MedChemExpress, HY-12947) was dissolved in DMSO and added to fly food at a final concentration of 35 µM. Larvae were grown directly on food supplemented with the DLK inhibitor. DMSO alone was used as a control (RUSSO AND DIANTONIO 2019).

### Immunohistochemistry and Imaging

Third instar wandering larvae were dissected in cold, calcium-free HL3 saline (pH 7.2) containing 70 mM NaCl, 5 mM KCl, 20 mM MgCl_2_, 10 mM NaHCO_3_, 5 mM Trehalose, 115 mM sucrose, and 5 mM HEPES and fixed in 4% paraformaldehyde for 30 min. Fillets were washed in PBS containing 0.2% Triton X-100, blocked for 1 hour in 5% bovine serum albumin (BSA), and then incubated overnight at 4 °C with the primary antibody. The primary antibodies used in the study are mouse anti-CSP (1:50) (DSHB, Iowa city, USA), mouse anti-NC82 (1:100) (DSHB), rat anti-Elav (1:200) (DSHB), mouse anti-β-galactosidase (1:200) (DSHB), mouse anti-mCherry (#M11217, Invitrogen), rabbit anti-p-eIF2α (#3597, CST), rabbit anti-Wallenda (1:100, (COLLINS *et al*. 2006)), The pMad antibody was a gift from Peter ten Dijke and used at 1:1000 dilution (PERSSON *et al*. 1998), guinea pig anti-Rab5 (1:500) (MOTTOLA *et al*. 2010) was a gift from Marino Zerial (Max Planck Institute, Germany), rabbit anti-Rab7 (1:500) and rabbit anti-Rab11 (1:500); (TANAKA AND NAKAMURA 2008; WEST *et al*. 2015) antibodies were a gift from Tsubasa Tanaka (RIKEN Center for Developmental Biology, Japan). Rabbit anti-Caprin (1:200) was a generous gift from Ophelia Papoulas (University of Texas, (PAPOULAS *et al*. 2010). The secondary antibodies conjugated to Alexa Fluor 488, Alexa Fluor 568, and Alexa Fluor 633 (Invitrogen, ThermoFisher Scientific, Waltham, MA, USA) were used at a 1:800 dilution. The Alexa Fluor 488- or Rhodamine-conjugated anti-HRP (Jackson ImmunoResearch, Pennsylvania, USA) was used at a 1:800 dilution.

For Wnd staining, larvae were dissected in ice-chilled 1X PBS and fixed in 3.7% formalin on ice for one hour. Larval fillets were blocked with 5% BSA for 1 hour, followed by overnight incubation with primary antibodies anti-Wnd and anti-NC82 at 4°C (COLLINS *et al*. 2006). For GFP-mCherry-Atg8a staining, the larvae were dissected in cold, calcium-free HL3 saline and fixed for 15 minutes at room temperature in 4% PFA in PBS (pH 7.2). Larval fillets were mounted in VECTASHIELD (Vector Laboratories, California, USA), and images were captured using a laser-scanning confocal microscope (FV3000, Olympus Corporation, Tokyo, Japan).

### Hexanediol treatment

Third instar *Drosophila* larvae expressing GFP-Hiw were dissected in cold Schneider’s *Drosophila* Medium (Gibco). Following dissection, preparations were gently washed with fresh Schneider’s medium to remove residual debris. To assess the solubility properties of accumulated Hiw, preparations were incubated for 2 minutes at room temperature in Schneider’s medium containing 1,6-hexanediol (1,6-HD; Sigma-Aldrich, #H6703) or 2,5-HD (Sigma-Aldrich, #H11904) at final concentrations of 0%, 1%, 5%, or 10%. The 0% condition consisted of Schneider’s medium alone (LIU *et al*. 2020). Following treatment, samples were immediately fixed in 4% paraformaldehyde in PBS for 30 minutes at room temperature. Fixed tissues were then washed and processed for immunostaining. No antibody amplification was used for GFP. The drug-containing medium was replaced with normal Schneider’s medium for 2 hr for recovery before fixation.

### Western Blotting

Third-instar larval *Drosophila* brains were homogenized in 1× SDS lysis buffer (50 mM Tris-Cl, pH 6.8; 25 mM KCl; 2 mM EDTA; 0.3 M sucrose; 2% SDS). The protein concentration was determined using the bicinchoninic acid (BCA) protein assay. Homogenates were mixed with an equal volume of 2× Laemmli buffer (50 mM Tris-HCl, pH 6.8; 2% SDS; 2% β-mercaptoethanol; 0.1% bromophenol blue; 10% glycerol), and 30 μg of protein was loaded onto a 5% SDS-PAGE gel. Proteins were transferred to a Hybond-LFP PVDF membrane (GE Healthcare, Illinois, USA). Membranes were blocked in 5% skimmed milk in 1× Tris-buffered saline containing 0.2% Tween-20 (0.2% TBST) for 1 hour at room temperature and incubated overnight at 4°C with primary antibodies: mouse anti-GFP (1:2000) and anti-Ran (1:2000, BD Biosciences, New Jersey, USA). After washing with 0.2% TBST, membranes were incubated with HRP-conjugated secondary antibody for 1 hour at room temperature. Signals were detected using the LI-COR Odyssey imaging system (LI-COR Biosciences, Lincoln, USA).

### Bafilomycin A1 and Chloroquine Treatment

To pharmacologically inhibit lysosomal function, third instar *Drosophila* larvae expressing GFP-Hiw were transferred to standard fly food containing either 200 nM Bafilomycin A1 (Baf) for 6 hours or 2.5 mg/ml Chloroquine (CQ) for 24 hours at 25°C. Stock solutions of both compounds were prepared in DMSO, and larvae fed on DMSO alone were served as controls. After the treatment, larvae were dissected in cold, calcium-free HL3 saline (pH 7.2) and proceeded with the immunocytochemistry protocol (ZIRIN *et al*. 2013; MAUVEZIN *et al*. 2015).

### Quantifications and Statistical Analysis

The NMJ 6/7 from the A2 hemisegment were captured to calculate the bouton number. All the CSP-positive boutons were counted manually in ImageJ/Fiji software (National Institutes of Health, Bethesda, USA). For bouton area quantification, all type-II boutons from muscle 6/7 at the A2 hemisegment were used, and the bouton area was calculated by drawing a freehand sketch around CSP-positive boutons using ImageJ; the average bouton area was represented in the graphs. Similarly, synaptic area was also manually quantified by a freehand sketch around the HRP-positive NMJ area. The total number of branches was manually counted in ImageJ.

Images were captured using a plan apochromat 60×, 1.42 NA objective for fluorescence intensity quantification. All control and experimental fillets were processed similarly for each experimental set, and fluorescence images were captured under identical settings. For GFP-atg8a levels and GFP-Hiw levels at NMJ, the average GFP intensity was normalized to the corresponding HRP intensity. The average fluorescence intensity of GFP was normalized to that of mCherry for the analysis of autophagy flux. The Hiw-GFP, GFP, TRE-DsRed, puc-LacZ, and endogenous mCherry-tagged Wnd levels were normalized to the corresponding Elav intensity in the VNC. The NC82-positive area was marked as the ROI, and the mean intensity for Wnd (antibody staining) was measured and normalized to the corresponding NC82 intensity.

To quantify the GFP-Hiw puncta number and area, z-projections of confocal images of third-instar larval VNCs were captured and stacked. The images were thresholded, and the Analyze Particles function in ImageJ was used to quantify the number of puncta and their size. For average puncta area, the mean area of all quantified puncta within each VNC was calculated, with each VNC represented as a single data point. For puncta number, only puncta with an area greater than 0.5 µm² were quantified using the Analyze Particles function in ImageJ.

The density of Western blot bands was quantified using ImageJ software. For multiple comparisons, one-way ANOVA followed by Tukey’s or Student’s t-tests was used. GraphPad Prism 8 (GraphPad Software Inc., California, USA) was used to plot all the graphs. Error bars in all the histograms represent SEM. *p < 0.05, **p < 0.01, ***p < 0.001, ns; not significant.

## Aknowledgments

We thank Dr. Avital Rodal for sharing *Rab11* mutants, Dr. Mohit Prasad for sharing the *puc-lacZ* fly lines, and Dr. Guang-Chao Chen for the *atg1* mutant lines. We also thank Dr. Aaron DiAntonio and Dr. Catherine Collins for generously sharing the Wallenda antibody. We thank Dr. Catherine Collins for sharing the endogenously tagged Wnd-mCherry fly line, Dr. Tsubasa Tanaka for Rab7 and Rab11 antibodies, Dr. Marino Zerial for Rab5 antibodies, Dr. Ophelia Papoulas for sharing the Caprin antibody; the BDSC for the fly stocks; DSHB, the University of Iowa, for monoclonal antibodies; and Varun Chaudhary, Sunando Datta, and Catherine Collins for their helpful comments on this manuscript. We thank Sakshi and Shruti for helping with the Western blots and figure organization. We thank the DST-FIST program of the Government of India for providing a confocal microscope to IISER-Bhopal. This work was supported by the Science and Engineering Board (SERB) Project Grant CRG/2021/000599, the Government of India, and by intramural funds from the Indian Institute of Science Education and Research Bhopal to V.K.

## Data availability

All data related to this manuscript are contained within the article.

## Notes

### Competing Interest Statement

The authors have declared no competing interest.

### Summary of Updates

The key changes in the revised manuscript are: 1. Strengthened the mechanistic interpretation and tempered claims regarding Hiw accumulations: We have revised the manuscript to describe the Hiw structures as dynamic accumulations of Hiw-positive puncta rather than bona fide LLPS condensates, acknowledging the limitations of our FRAP and LLPS experiments. We have also moderated mechanistic claims concerning endocytosis-dependent Hiw localization and its potential contribution to Wnd/JNK activation, clearly distinguishing experimental findings from the proposed model and acknowledging alternative mechanisms. In view of our limitation in showing the Hiw puncta as bona fide LLPS condensates, we have changed the title of the manuscript to "E3 Ubiquitin Ligase Highwire/Phr1 Mediates Endocytic Control of JNK Signaling in Drosophila Neurons". 2. Expanded experimental validation and strengthened genetic evidence: We have incorporated additional experiments to validate key findings, including analysis of 1,6-hexanediol recovery in a time-course manner, and shits experiments with short-duration temperature shifts at restrictive temperatures to assess time-dependency of Hiw accumulation. We have also reverified the Wnd levels using endogenously mCherry-tagged Wnd and pharmacologically blocked Wnd activity with GNE-3511 to independently confirm the Wnd-dependent regulation of synaptic overgrowth in the endocytic mutants. Following the reviewer's suggestions, we have reverified the genetic data with homozygous atg1 and wit mutants, and performed p-Mad staining to assess BMP pathway activation. Furthermore, we have expanded quantification of synaptic growth parameters, improved image quality and antibody staining presentation for Wnd, and examined the reproducibility of the Hiw accumulation using additional neuronal drivers (D42-Gal4 and nSyb-Gal4). 3. Rewriting Results and Discussion: We have clarified normalization procedures, corrected terminology and figure labels, and expanded the quantitative analysis of synaptic architecture, including bouton number and size, synaptic area, branch number and length, and satellite boutons. We have also strengthened the Discussion by explicitly acknowledging experimental limitations and distinguishing established findings from unresolved mechanistic questions.

