## Supplemental data for "E3 Ubiquitin Ligase Highwire/Phr1 Mediates Endocytic Control of JNK Signaling in *Drosophila* Neurons"

### Supplemental figures

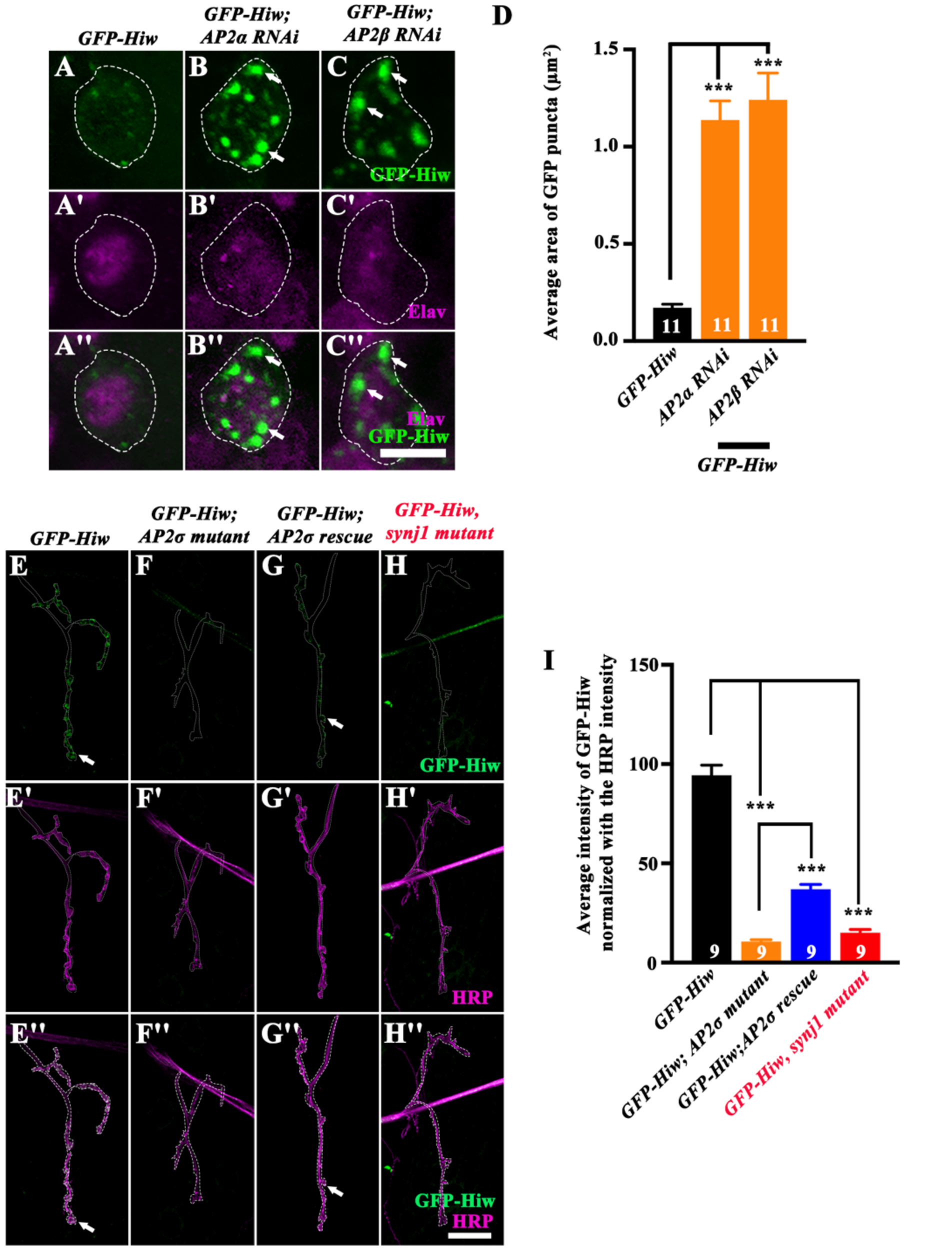


**Figure_1 - supplement 1**

**Figure 1 – figure supplement 1: Hiw trafficking is defective in endocytic mutants**

(**A-C''**) Confocal images of the neuronal cell body co-labeled for GFP-Hiw (green) and Elav (magenta) in control (*Elav-Gal4>UAS-GFP-Hiw*) (A-A''), *Elav-Gal4>UAS-GFP-Hiw; UAS-AP2α RNAi* (B-B''), and *Elav-Gal4>UAS-GFP-Hiw; UAS-AP2β RNAi* (C-C''). The scale bar in C'' represents 5 μm. Arrows point to the Hiw puncta in the cell body.

(**D**) Histogram showing the average area ($\mu m^2$) of GFP-Hiw puncta in control (0.17 ± 0.02), *Elav-Gal4>UAS-GFP-Hiw; UAS-AP2α RNAi* (1.14 ± 0.10), and *Elav-Gal4>UAS-GFP-Hiw; UAS-AP2β RNAi* (1.24 ± 0.14). At least 11 VNCs of each genotype were used for the quantifications. The error bar represents the standard error of the mean (SEM); the statistical analysis was done using one-way ANOVA followed by post hoc Tukey's test. ***p<0.001.

(**E-H''**) Confocal images of muscle 4 NMJ showing the synaptic levels of GFP-Hiw in *Elav-Gal4>UAS-GFP-Hiw* (E-E''), *Elav-Gal4>UAS-GFP-Hiw;* $AP2\sigma^{ang7}/AP2\sigma\text{-}^{KG}$ (F-F''), *Elav-Gal4>UAS-GFP-Hiw; UAS-AP2σ,* $AP2\sigma^{ang7}/AP2\sigma\text{-}^{KG}$ (G-G''), and *Elav-Gal4>UAS-GFP-Hiw,* $synj1^{1/2}$ (H-H'') immunolabeled for HRP (magenta) and GFP-Hiw (green). The scale bar in H'' represents 10 μm. Arrows point to the Hiw at the synaptic terminals.

(**I**) Histogram showing the synaptic levels of the GFP-Hiw normalized with the HRP at the larval NMJ of muscle 4 of control (*Elav-Gal4>UAS-GFP-Hiw*) (93.7 ± 5.22), *Elav-Gal4>UAS-GFP-Hiw;* $AP2\sigma^{ang7}/AP2\sigma\text{-}^{KG}$ (10.7 ± 0.92), *Elav-Gal4>UAS-GFP-Hiw; UAS-AP2σ,* $AP2\sigma^{ang7}/AP2\sigma\text{-}^{KG}$ (36.9 ± 2.43), and *Elav-Gal4>UAS-GFP-Hiw,* $synj1^{1/2}$ (15 ± 1.71). At least 9 NMJs from each genotype were used for quantification. The error bar represents the standard error of the mean (SEM); the statistical analysis was done using one-way ANOVA followed by post hoc Tukey's test. ***p<0.001.

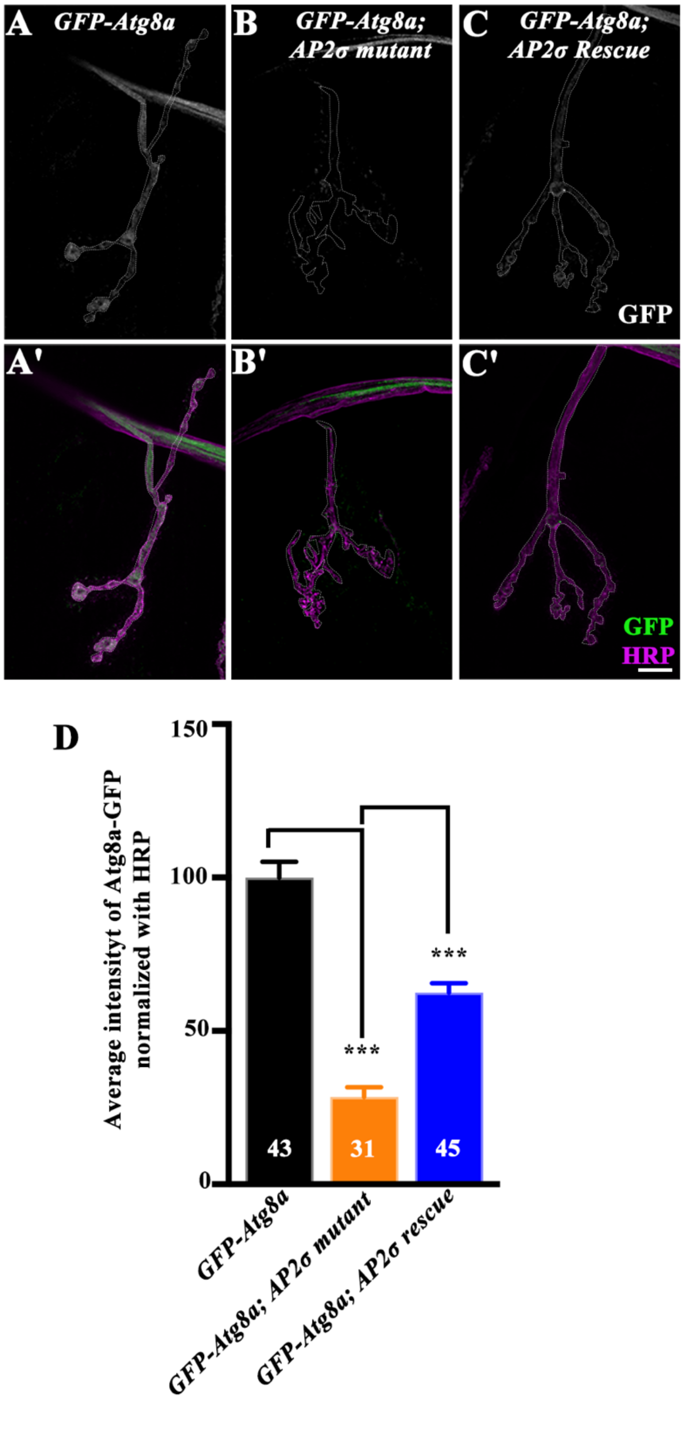


**Figure 1 - figure supplement 2**

**Figure 1 – figure supplement 2: Autophagic marker, Atg8a, is reduced in *AP2* mutant synapses**

(**A-C'**) Confocal images of muscle 4 NMJ co-labeled for GFP-Atg8a (green) and HRP (magenta) in control (*Elav-Gal4>UAS-GFP-Atg8a*) (A-A'), *Elav-Gal4>UAS-GFP-Atg8a;*

*AP2σ$^{ang7}$/AP2σ-$^{KG}$* (B-B'), *and Elav-Gal4>UAS-GFP-Atg8a; UAS-AP2σ, AP2σ$^{ang7}$/AP2σ-$^{KG}$* (C-C'). The scale bar in C' represents 10 µm for A-C'.

(**D**) Histogram showing the Atg8a-GFP levels at the muscle 4 NMJ of the control (*Elav-Gal4>UAS-GFP-Atg8a*) (100.0 ± 5.15)*, Elav-Gal4>UAS-GFP-Atg8a; AP2σ$^{ang7}$/AP2σ-$^{KG}$* (28.49 ± 3.18), *and Elav-Gal4>UAS-GFP-Atg8a; UAS-AP2σ, AP2σ$^{ang7}$/AP2σ-$^{KG}$* (62.43 ± 3.15). At least 31 NMJs from each genotype were used for quantification. The error bar represents the standard error of the mean (SEM); the statistical analysis was done using one-way ANOVA followed by post hoc Tukey's test. ***p<0.001.

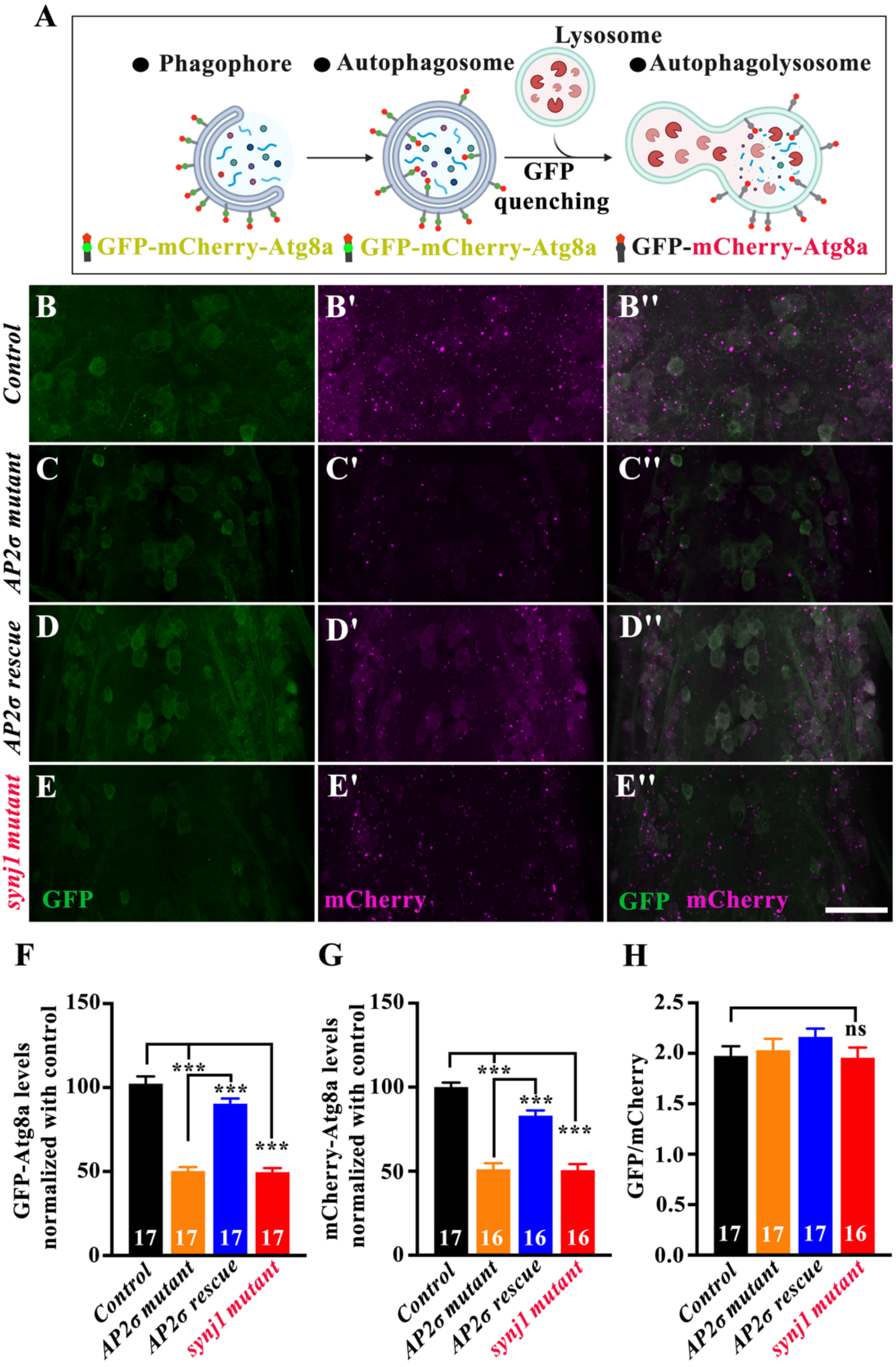


Figure 1 - figure supplement 3

**Figure 1 – figure supplement 3: Endocytosis regulates autophagosome formation in neurons**

(**A**) Cartoon representation showing trafficking of the GFP-mCherry-Atg8a during the autophagic process. In the early stages of autophagosome formation, both GFP and mCherry fluoresce, so the autophagosomes appear yellow due to the overlap of green and red fluorescence. During the later stages, the autophagosome fuses with the lysosome to form an autolysosome. The acidic environment of the lysosome quenches GFP but not mCherry, causing autolysosomes to appear red, indicating the completion of the autophagic process.

(**B-E''**) Confocal images of third instar larval ventral nerve cord co-labeled for GFP (green) and mCherry (magenta) in control (*Elav-Gal4>UAS-GFP-mCherry-Atg8a*) (B-B''), *Elav-Gal4>UAS-GFP-mCherry-Atg8a; AP2σ-*$^{KG}$*/AP2σ*$^{ang7}$ (C-C''), *Elav-Gal4>UAS-GFP-mCherry-Atg8a; UAS-AP2σ, AP2σ-*$^{KG}$*/AP2σ*$^{ang7}$ (D-D''), and *Elav-Gal4>UAS-GFP-mCherry-Atg8a; synj1*$^{1/2}$ (E-E''). The scale bar in E'' represents 40 µm.

(**F**) Histogram showing GFP-Atg8a levels in the neuronal cell body of control (*Elav-Gal4>UAS-GFP-mCherry-Atg8a*) (102.3 ± 4.30), *Elav-Gal4>UAS-GFP-mCherry-Atg8a; AP2σ-*$^{KG}$*/AP2σ*$^{ang7}$ (50.26 ± 2.35), *Elav-Gal4>UAS-GFP-mCherry-Atg8a; UAS-AP2σ, AP2σ-*$^{KG}$*/AP2σ*$^{ang7}$ (90.37 ± 3.07), and *Elav-Gal4>UAS-GFP-mCherry-Atg8a; synj1*$^{1/2}$ (49.59 ± 2.47). At least 17 VNCs of each genotype were used for the quantification.

(**G**) Histogram showing mCherry-Atg8a levels in the neuronal cell body of control (*Elav-Gal4>UAS-GFP-mCherry-Atg8a*) (100.0 ± 2.73), *Elav-Gal4>UAS-GFP-mCherry-Atg8a; AP2σ-*$^{KG}$*/AP2σ*$^{ang7}$ (51.09 ± 3.64), *Elav-Gal4>UAS-GFP-mCherry-Atg8a; UAS-AP2σ, AP2σ-*$^{KG}$*/AP2σ*$^{ang7}$ (83.05 ± 3.08), and *Elav-Gal4>UAS-GFP-mCherry-Atg8a; synj1*$^{1/2}$ (50.63 ± 3.58). At least 16 VNCs of each genotype were used for the quantification.

(**H**) Histogram showing GFP to mCherry ratio in the neuronal cell body of control (*Elav-Gal4>UAS-GFP-mCherry-Atg8a*) (1.97 ± 0.09), *Elav-Gal4>UAS-GFP-mCherry-Atg8a; AP2σ-*$^{KG}$*/AP2σ*$^{ang7}$ (2.03 ± 0.11), *Elav-Gal4>UAS-GFP-mCherry-Atg8a; UAS-AP2σ, AP2σ-*$^{KG}$*/AP2σ*$^{ang7}$ (2.16 ± 0.08), and *Elav-Gal4>UAS-GFP-mCherry-Atg8a; synj1*$^{1/2}$ (1.95 ± 0.10).

At least 16 VNCs of each genotype were used for the quantification. The error bars in F, G, and H represent the standard error of the mean (SEM); the statistical analysis was performed using one-way ANOVA followed by Tukey's post hoc test. ***$p<0.001$, ns: not significant.

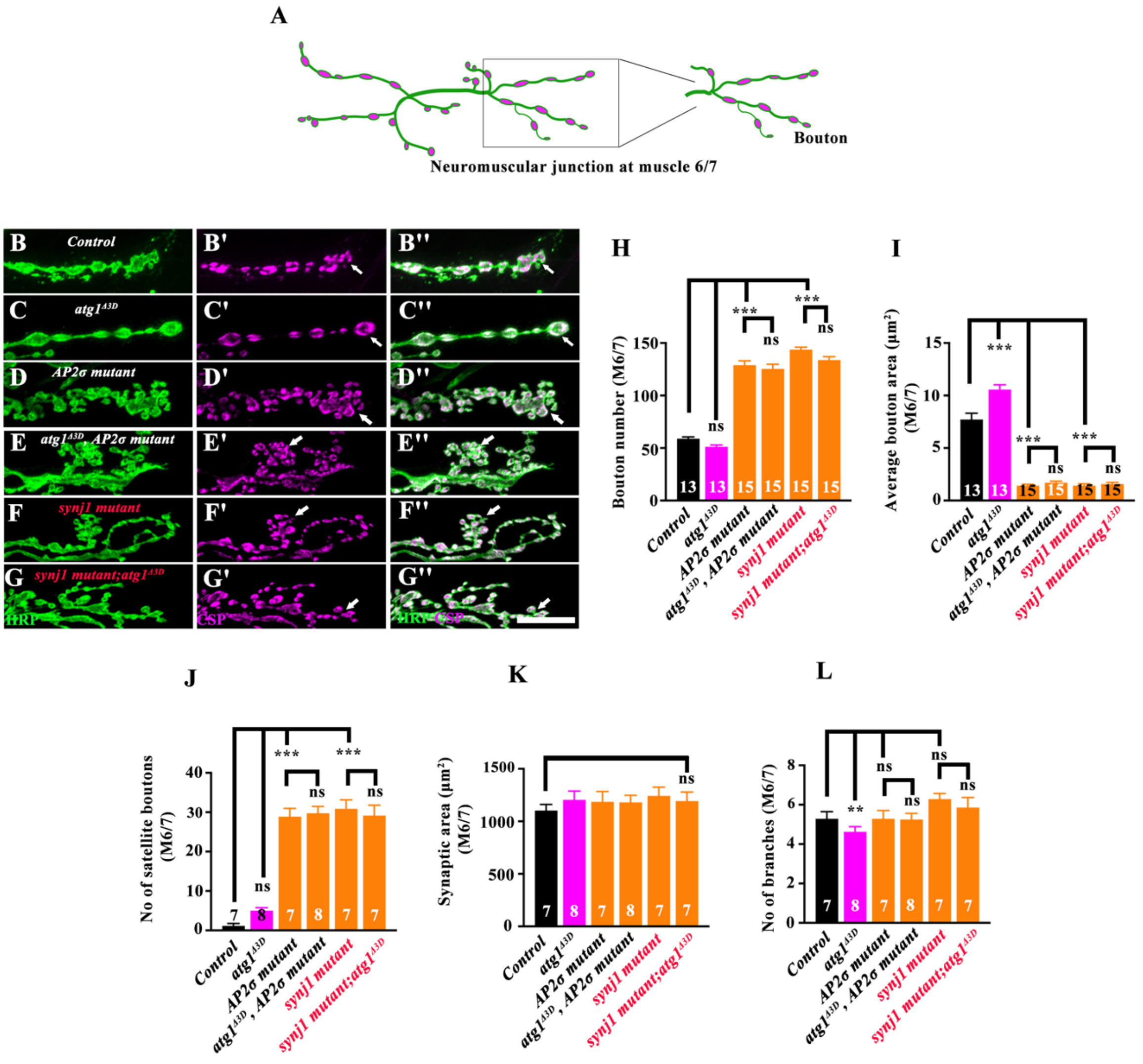


**Figure 1 - figure supplement 4**

**Figure 1 – figure supplement 4: Autophagy does not influence the NMJ phenotypes in the AP2 mutants.**

(**A**) Schematic representation of the third instar larval NMJ at muscle 6/7. The magnified view shows the terminal synaptic bouton phenotypes.

(**B-G''**) Confocal images of muscle 6/7 NMJ showing the synaptic growth in control (B-B''), *atg1*$^{Δ3D}$ (C-C''), *AP2σ*$^{KG}$/*AP2σ*$^{ang7}$ (D-D''), *atg1*$^{Δ3D}$, *AP2σ*$^{KG}$/*AP2σ*$^{ang7}$ (E-E''), *synj1*$^{1/2}$ (F-F''), and *synj1*$^{1/2}$; *atg1*$^{Δ3D}$ (G-G'') immunolabeled for CSP (magenta) and HRP (green). The scale bar in G'' represents 10 μm. The arrows indicate changes in NMJ morphology.

(**H**) Histogram showing the bouton number from muscle 6/7 NMJ at A2 hemisegment for the control (58.9 ± 1.72), *atg1*$^{\Delta3D}$ (51.2 ± 1.67), *AP2σ*$^{KG}$/*AP2σ*$^{ang7}$ (129 ± 4.07), *atg1*$^{\Delta3D}$, *AP2σ*$^{KG}$/*AP2σ*$^{ang7}$ (125 ± 4.43), *synj1*$^{1/2}$ (144 ± 2.28), and *synj1*$^{1/2}$; *atg1*$^{\Delta3D}$ (134 ± 3.27). At least 13 NMJs from each genotype were used for quantification.

(**I**) Histogram showing the average bouton area (µm$^2$) from muscle 6/7 NMJ at A2 hemisegment for the control (7.71 ± 0.6), *atg1*$^{\Delta3D}$ (10.6 ± 0.452), *AP2σ*$^{KG}$/*AP2σ*$^{ang7}$ (1.41 ± 0.13), *atg1*$^{\Delta3D}$, *AP2σ*$^{KG}$/*AP2σ*$^{ang7}$ (1.7 ± 0.15), *synj1*$^{1/2}$ (1.41 ± 0.18), and *synj1*$^{1/2}$; *atg1*$^{\Delta3D}$ (1.57 ± 0.15). At least 13 NMJs from each genotype were used for quantification.

(**J**) Histogram showing the number of satellite boutons in muscle 6/7 NMJ at A2 hemisegment for the control (1.14 ± 0.6), *atg1*$^{\Delta3D}$ (5 ± 0.75), *AP2σ*$^{KG}$/*AP2σ*$^{ang7}$ (28.9 ± 2.12), *atg1*$^{\Delta3D}$, *AP2σ*$^{KG}$/*AP2σ*$^{ang7}$ (29.8 ± 1.73), *synj1*$^{1/2}$ (30.9 ± 2.29), and *synj1*$^{1/2}$; *atg1*$^{\Delta3D}$ (29.1 ± 2.65). At least 7 NMJs from each genotype were used for quantification.

(**K**) Histogram showing the synaptic area (µm$^2$) of muscle 6/7 NMJ at A2 hemisegment for the control (1102 ± 58.2), *atg1*$^{\Delta3D}$ (1205 ± 83.4), *AP2σ*$^{KG}$/*AP2σ*$^{ang7}$ (1185 ± 98.4), *atg1*$^{\Delta3D}$, *AP2σ*$^{KG}$/*AP2σ*$^{ang7}$ (1181 ± 67.4), *synj1*$^{1/2}$ (1242 ± 83.5), and *synj1*$^{1/2}$; *atg1*$^{\Delta3D}$ (1193 ± 86.2). At least 7 NMJs of each genotype were used for quantification.

(**L**) Histogram showing the number of branches in muscle 6/7 NMJ at A2 hemisegment for the control (5.29 ± 0.36), *atg1*$^{\Delta3D}$ (4.63 ± 0.26), *AP2σ*$^{KG}$/*AP2σ*$^{ang7}$ (5.29 ± 0.42), *atg1*$^{\Delta3D}$, *AP2σ*$^{KG}$/*AP2σ*$^{ang7}$ (5.25 ± 0.31), *synj1*$^{1/2}$ (6.29 ± 0.28), and *synj1*$^{1/2}$; *atg1*$^{\Delta3D}$ (5.86 ± 0.51). At least 7 NMJs from each genotype were used for quantification. The error bars in H, I, J, K, and L represent the standard error of the mean (SEM); the statistical analysis was performed using one-way ANOVA followed by Tukey's post hoc test. ***$p<0.001$; **$p<0.01$, ns: not significant.

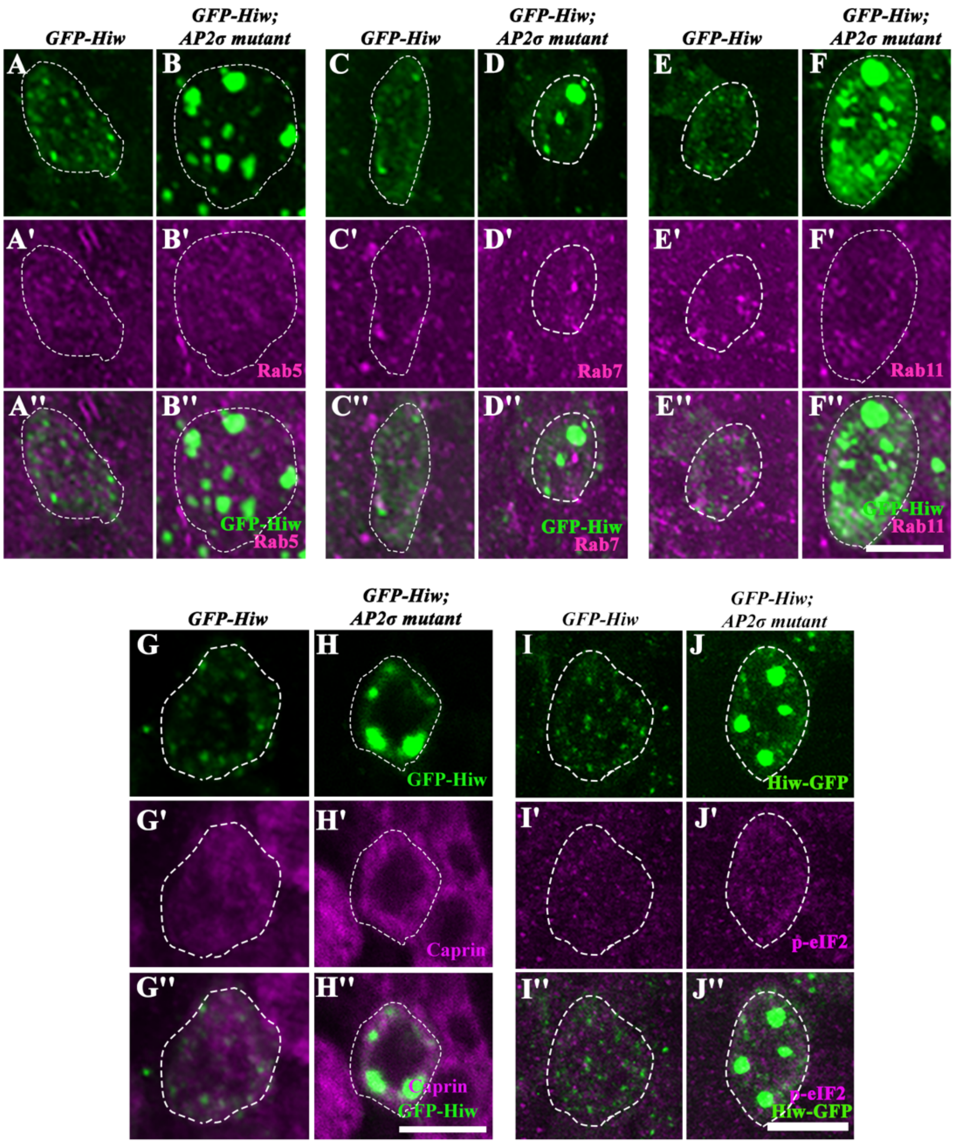


**Figure 2 - figure supplement 1**

**Figure 2 – figure supplement 1: Hiw accumulations do not colocalize with endosomal or stress granule markers**

(**A- J''**) Confocal images of third instar larval neuronal cell bodies in control (*Elav-Gal4>UAS-GFP-Hiw*) and *Elav-Gal4>UAS-GFP-Hiw; AP2σ*$^{ang7}$*/AP2σ-*$^{KG}$ showing colocalization between the GFP-Hiw (green) and Rab5 (magenta) (A-B''), GFP-Hiw (green) and Rab7 (magenta) (C-D''), GFP-Hiw (green) and Rab11 (magenta) (E-F''), GFP-Hiw (green) and Caprin (magenta) (G-H''), and GFP-Hiw (green) and p-eIF2α (magenta) (I-J''). The scale bar in F'' represents 5 μm for images A-F'', the scale bar in H'' represents 5 μm for G-H'', and the scale bar in J'' represents 5 μm for I-J''.

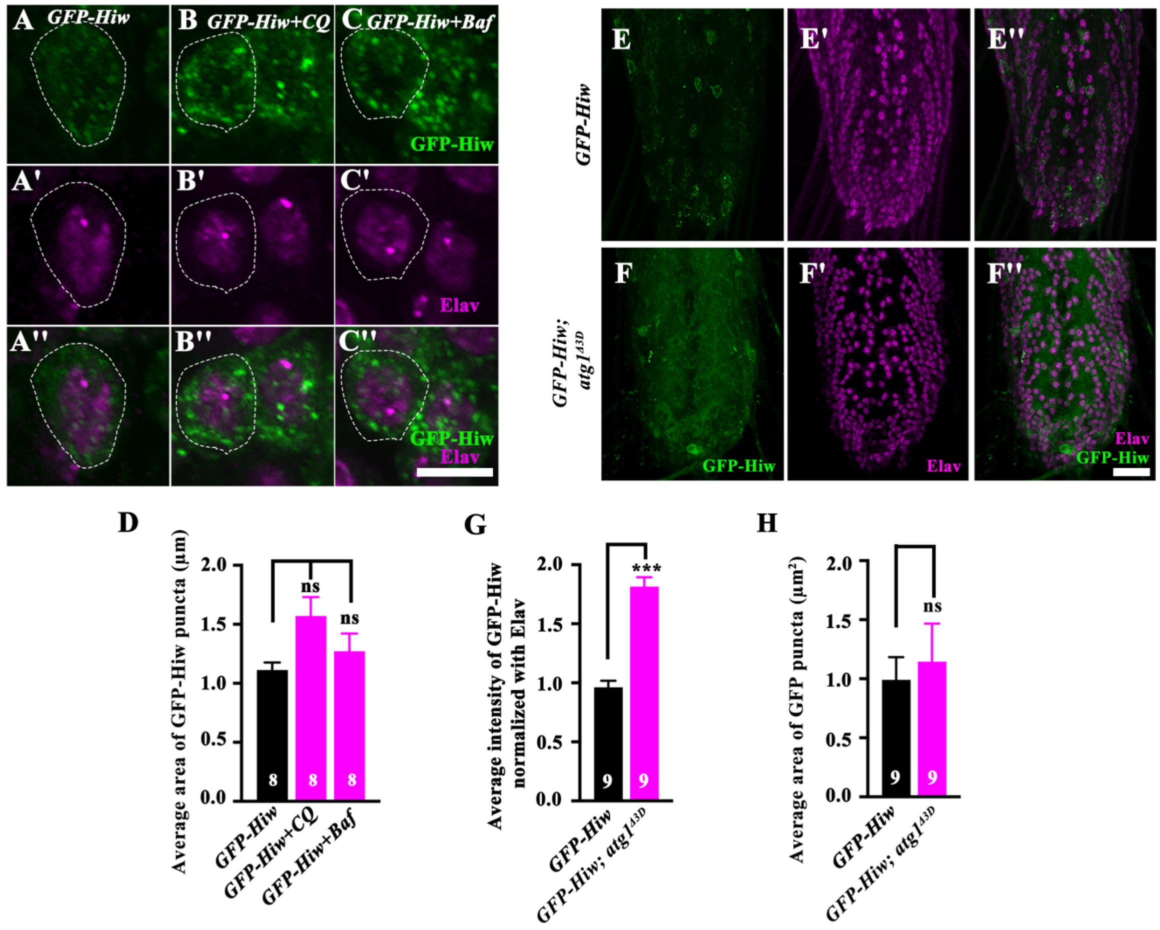


**Figure 2 - figure supplement 2**

**Figure 2 – figure supplement 2: Hiw accumulations are not due to the reduced levels of autophagy in the endocytic mutants.**

(**A-C''**) Confocal images of the neuronal cell body co-labeled for GFP-Hiw (green) and Elav (magenta) in the control (*Elav-Gal4>UAS-GFP-Hiw*) (A-A''), *Elav-Gal4>UAS-GFP-Hiw* + Chloroquine (B-B''), and *Elav-Gal4>UAS-GFP-Hiw* + Bafilomycin A1 (C-C''). The scale bar in C'' represents 5 µm.

(**D**) Histogram showing the average area (µm$^2$) of the GFP puncta in the neuronal cell body of the control (*Elav-Gal4>UAS-GFP-Hiw*) (0.11 ± 0.01), *Elav-Gal4>UAS-GFP-Hiw* + Chloroquine (0.15 ± 0.02), and *Elav-Gal4>UAS-GFP-Hiw* + Bafilomycin A1 (0.13 ± 0.01). At least 8 VNCs from each genotype were used for quantification.

(**E-F''**) Confocal images of the third-instar larval VNC co-labeled for GFP-Hiw (green) and Elav (magenta) in the control (*Elav-Gal4>UAS-GFP-Hiw*) (E-E''), *Elav-Gal4>UAS-GFP-Hiw; atg1*$^{\Delta 3D}$ (F-F''). The scale bar in F'' represents 10 µm.

(**G**) Histogram showing the average intensity of GFP-Hiw normalized with Elav in control (*Elav-Gal4>UAS-GFP-Hiw*) (0.96 ± 0.05), *Elav-Gal4>UAS-GFP-Hiw; atg1*$^{\Delta 3D}$ (1.82 ± 0.08). At least 9 VNCs of each genotype were used for quantification.

(**H**) Histogram showing the average area ($\mu m^2$) of the GFP puncta in the neuronal cell body of the control (*Elav-Gal4>UAS-GFP-Hiw*) (0.198 ± 0.04) and *Elav-Gal4>UAS-GFP-Hiw; atg1*$^{\Delta 3D}$ (0.23 ± 0.06). At least 12 VNCs of each genotype were used for quantification. The error bars in D, G, and H represent the standard error of the mean (SEM); the statistical analysis was done using Student's t-test (G and H). The statistical analysis was done using one-way ANOVA followed by post hoc Tukey's test (D). ***$p<0.001$, ns: not significant.

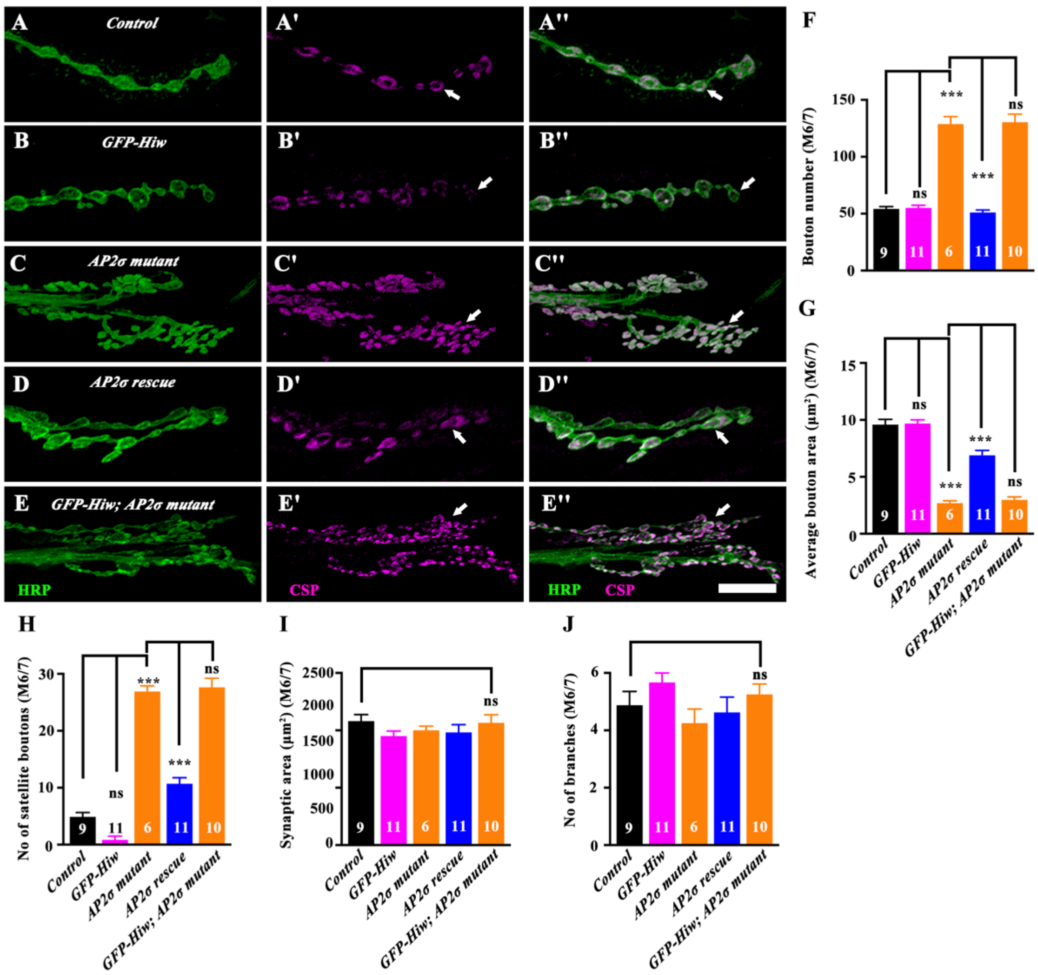


Figure 4 - figure supplement 1

**Figure 4 – figure supplement 1: Neuronal expression of Hiw does not suppress the synaptic overgrowth in *σ2-adaptin* mutants.**

(**A-E''**) Confocal images of muscle 6/7 NMJ immunolabeled for CSP (magenta) and HRP (green) to show the synaptic growth phenotypes in the control (A-A''), *Elav-Gal4>UAS-GFP-*Hiw (B-B''), *AP2σ*$^{ang7}$*/AP2σ-*$^{KG}$ (C-C''), *Elav-Gal4>UAS-AP2σ, AP2σ*$^{ang7}$*/AP2σ-*$^{KG}$ (D-D''), and *Elav-Gal4>UAS-GFP-Hiw; AP2σ*$^{ang7}$*/AP2σ-*$^{KG}$ (E-E''). The scale bar in E'' represents 10 µm. The arrows indicate changes in NMJ morphology.

(**F**) Histogram showing the bouton number at muscle 6/7 NMJ of A2 hemisegment for control (54.2 ± 1.98), *Elav-Gal4>UAS-GFP-Hiw* (55.0 ± 2.39), *AP2σ*$^{ang7}$*/AP2σ-*$^{KG}$ (129 ± 6.55), *Elav-*

*Gal4>UAS-AP2σ,* $AP2\sigma^{ang7}/AP2\sigma\text{-}^{KG}$ (51 ± 2.22), and *Elav-Gal4>UAS-GFP-Hiw;* $AP2\sigma^{ang7}/AP2\sigma\text{-}^{KG}$ (130 ± 7.06). At least 6 NMJs of each genotype were used for quantification.

(**G**) Histograms showing the average bouton area ($\mu m^2$) at muscle 6/7 NMJ of A2 hemisegment for control (9.59 ± 0.46), *Elav-Gal4>UAS-GFP-Hiw* (9.68 ± 0.33), $AP2\sigma^{ang7}/AP2\sigma\text{-}^{KG}$ (2.69 ± 0.21), *Elav-Gal4>UAS-AP2σ,* $AP2\sigma^{ang7}/AP2\sigma\text{-}^{KG}$ (6.89 ± 0.43), and *Elav-Gal4>UAS-GFP-Hiw;* $AP2\sigma^{ang7}/AP2\sigma\text{-}^{KG}$ (2.97 ± 0.27). At least 6 NMJs of each genotype were used for quantification.

(**H**) Histograms showing the number of satellite boutons at muscle 6/7 NMJ of A2 hemisegment for control (4.88 ± 0.76), *Elav-Gal4>UAS-GFP-Hiw* (0.83 ± 0.65), $AP2\sigma^{ang7}/AP2\sigma\text{-}^{KG}$ (26.9 ± 1.01), *Elav-Gal4>UAS-AP2σ,* $AP2\sigma^{ang7}/AP2\sigma\text{-}^{KG}$ (10.7 ± 1.07), and *Elav-Gal4>UAS-GFP-Hiw;* $AP2\sigma^{ang7}/AP2\sigma\text{-}^{KG}$ (27.6 ± 1.59). At least 6 NMJs of each genotype were used for quantification.

(**I**) Histograms showing the synaptic area ($\mu m^2$) at muscle 6/7 NMJ of A2 hemisegment for control (1080 ± 57.3), *Elav-Gal4>UAS-GFP-Hiw* (951 ± 43), $AP2\sigma^{ang7}/AP2\sigma\text{-}^{KG}$ (1000 ± 36.1), *Elav-Gal4>UAS-AP2σ,* $AP2\sigma^{ang7}/AP2\sigma\text{-}^{KG}$ (982 ± 68), and *Elav-Gal4>UAS-GFP-Hiw;* $AP2\sigma^{ang7}/AP2\sigma\text{-}^{KG}$ (1065 ± 70.1). At least 6 NMJs of each genotype were used for quantification.

(**J**) Histograms showing the number of branches at muscle 6/7 NMJ of A2 hemisegment for control (4.88 ± 0.48), *Elav-Gal4>UAS-GFP-Hiw* (5.67 ± 0.33), $AP2\sigma^{ang7}/AP2\sigma\text{-}^{KG}$ (4.25 ± 0.50), *Elav-Gal4>UAS-AP2σ,* $AP2\sigma^{ang7}/AP2\sigma\text{-}^{KG}$ (4.63 ± 0.53), and *Elav-Gal4>UAS-GFP-Hiw;* $AP2\sigma^{ang7}/AP2\sigma\text{-}^{KG}$ (5.25 ± 0.36). At least 6 NMJs of each genotype were used for quantification. The error bars in F, G, H, I, and J represent the standard error of the mean (SEM); the statistical analysis was performed using one-way ANOVA followed by Tukey's post hoc test. ***p<0.001, ns: not significant.

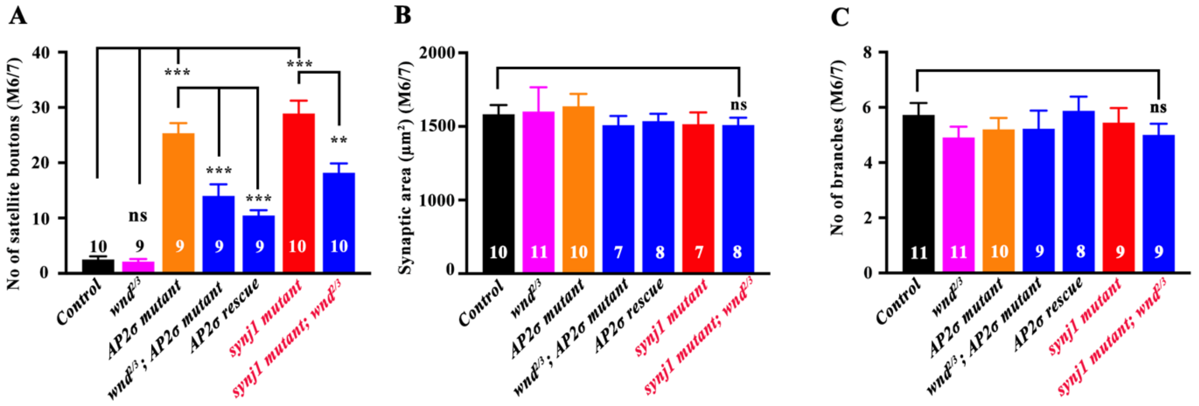

**Figure 4 - figure supplement 2**

**Figure 4-2: Redcution in Wnd levels rescues synaptic morphological defects of endocytic mutants**

(**A**) Histogram showing the number of satellite boutons at muscle 6/7 NMJ of A2 hemisegment for control (2.5 ± 0.58), *wnd*$^{2/3}$ (2.11 ± 0.48), *AP2σ*$^{ang7}$*/AP2σ-*$^{KG}$ (25.3 ± 1.83), *wnd*$^{2/3}$*, AP2σ*$^{ang7}$*/AP2σ-*$^{KG}$ (10.4 ± 0.96), *Elav-Gal4>UAS-AP2σ, AP2σ*$^{ang7}$*/AP2σ-*$^{KG}$ (14 ± 2.09), *synj*$^{1/2}$ (28.9 ± 2.33), and *synj*$^{1/2}$*; wnd*$^{2/3}$ (18.2 ± 1.67). At least 9 NMJs of each genotype were used for quantification.

(**B**) Histogram showing the synaptic area (μm$^2$) at muscle 6/7 NMJ of A2 hemisegment for control (1083 ± 62.5), *wnd*$^{2/3}$ (1101 ± 165), *AP2σ*$^{ang7}$*/AP2σ-*$^{KG}$ (1137 ± 83.8), *wnd*$^{2/3}$*, AP2σ*$^{ang7}$*/AP2σ-*$^{KG}$ (1037 ± 49.6), *Elav-Gal4>UAS-AP2σ, AP2σ*$^{ang7}$*/AP2σ-*$^{KG}$ (1009 ± 62.1), *synj*$^{1/2}$ (1015 ± 80.5), and *synj*$^{1/2}$*; wnd*$^{2/3}$ (1009 ± 50). At least 7 NMJs of each genotype were used for quantification.

(**C**) Histogram showing the number of branches at muscle 6/7 NMJ of A2 hemisegment for control (5.73 ± 0.43), $wnd^{2/3}$ (4.91 ± 0.32), $AP2\sigma^{ang7}/AP2\sigma\text{-}^{KG}$ (5.2 ± 0.41), $wnd^{2/3}$, $AP2\sigma^{ang7}/AP2\sigma\text{-}^{KG}$ (5.88 ± 0.51), *Elav-Gal4>UAS-AP2σ* $AP2\sigma^{ang7}/AP2\sigma\text{-}^{KG}$ (5.22 ± 0.66), $synj^{1/2}$ (5.44 ± 0.53), and $synj^{1/2}$*;* $wnd^{2/3}$ (5 ± 0.41). At least 8 NMJs of each genotype were used for quantification. The error bars in A, B, and C represent the standard error of the mean (SEM); the statistical analysis was done using one-way ANOVA followed by post hoc Tukey's test. ***p<0.001; **p<0.01; ns: not significant.

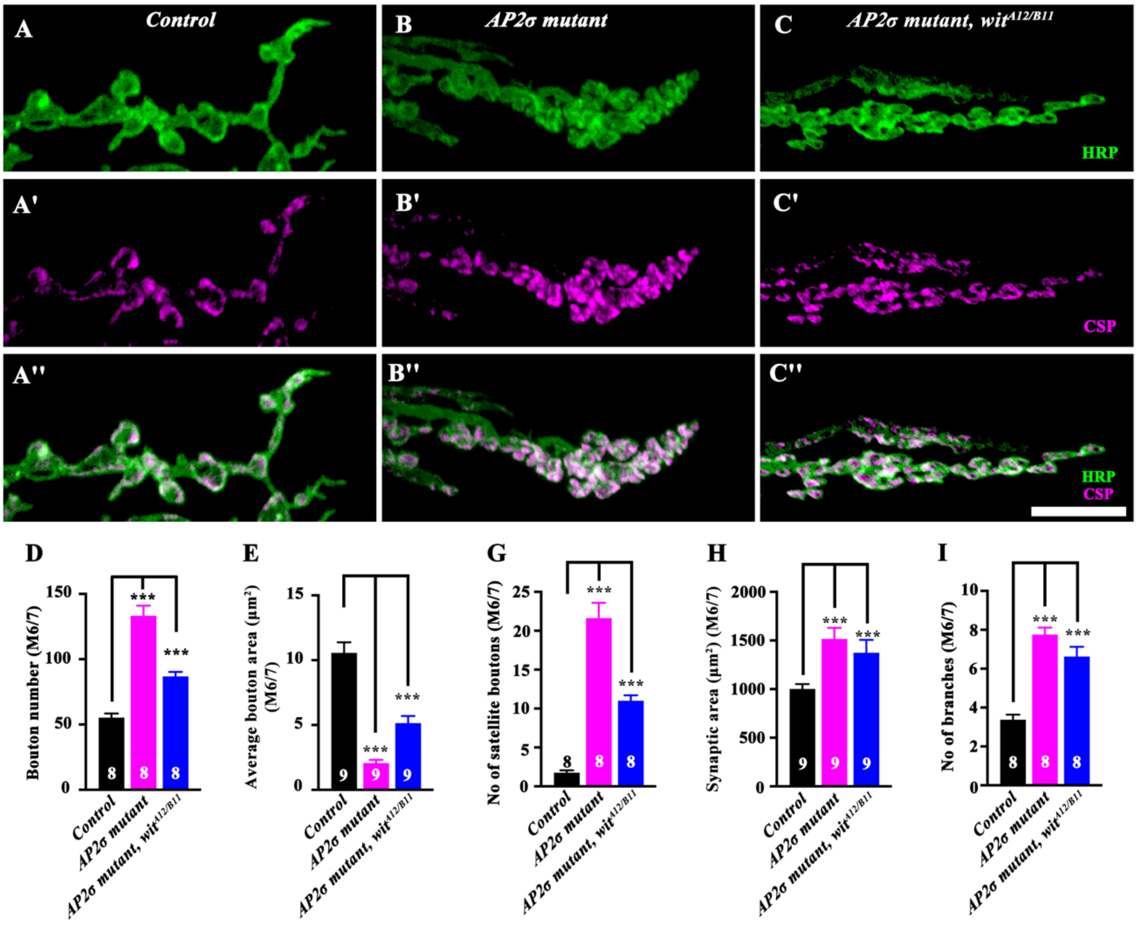


Figure 6 - figure supplement 1

**Figure 6 - figure supplement 1: Reduction in BMP signaling alone only partially rescues NMJ overgrowth in endocytic mutants**

(**A-C''**) Confocal images of muscle 6/7 NMJ immunolabeled for CSP (magenta) and HRP (green) to show the synaptic growth phenotypes in the control (A-A''), *AP2σ$^{ang7}$/AP2σ-$^{KG}$* (B-B''), and *AP2σ$^{ang7}$/AP2σ-$^{KG}$, wit$^{A12/B11}$* (C-C''). The scale bar in C'' represents 10 µm. The arrows indicate changes in NMJ morphology.

(**D**) Histograms showing the bouton number at muscle 6/7 NMJ of A2 hemisegment for control (55.3 ± 3.23), *AP2σ*$^{ang7}$*/AP2σ-*$^{KG}$ (133 ± 7.80), *AP2σ*$^{ang7}$*/AP2σ-*$^{KG}$*, wit*$^{A12/B11}$(86.9 ± 3.40). At least 8 NMJs of each genotype were used for quantification.

(**E**) Histograms showing the average bouton area ($\mu m^2$) at muscle 6/7 NMJ of A2 hemisegment for control (10.6 ± 0.82), *AP2σ*$^{ang7}$*/AP2σ-*$^{KG}$ (2.06 ± 0.25), *AP2σ*$^{ang7}$*/AP2σ-*$^{KG}$*, wit*$^{A12/B11}$(5.14 ± 0.55). At least 9 NMJs of each genotype were used for quantification.

(**F**) Histograms showing the number of satellite boutons at the muscle 6/7 NMJ of A2 hemisegment for control (1.75 ± 0.31), *AP2σ*$^{ang7}$*/AP2σ-*$^{KG}$ (21.6 ± 1.96), *AP2σ*$^{ang7}$*/AP2σ-*$^{KG}$*, wit*$^{A12/B11}$(11 ± 0.71). At least 8 NMJs of each genotype were used for quantification.

(**G**) Histograms showing the synaptic area ($\mu m^2$) at muscle 6/7 NMJ of A2 hemisegment for control (1001 ± 50.1), *AP2σ*$^{ang7}$*/AP2σ-*$^{KG}$ (1516 ± 114), *AP2σ*$^{ang7}$*/AP2σ-*$^{KG}$*, wit*$^{A12/B11}$(1374 ± 132). At least 9 NMJs of each genotype were used for quantification

(**H**) Histograms showing the number of branches at muscle 6/7 NMJ of A2 hemisegment for control (3.38 ± 0.26), *AP2σ*$^{ang7}$*/AP2σ-*$^{KG}$ (7.75 ± 0.36), *AP2σ*$^{ang7}$*/AP2σ-*$^{KG}$*, wit*$^{A12/B11}$(6.63 ± 0.50). At least 8 NMJs of each genotype were used for quantification. The error bars in D, E, F, G, and H represent the standard error of the mean (SEM); the statistical analysis was performed using one-way ANOVA followed by Tukey's post hoc test. ***p<0.001.

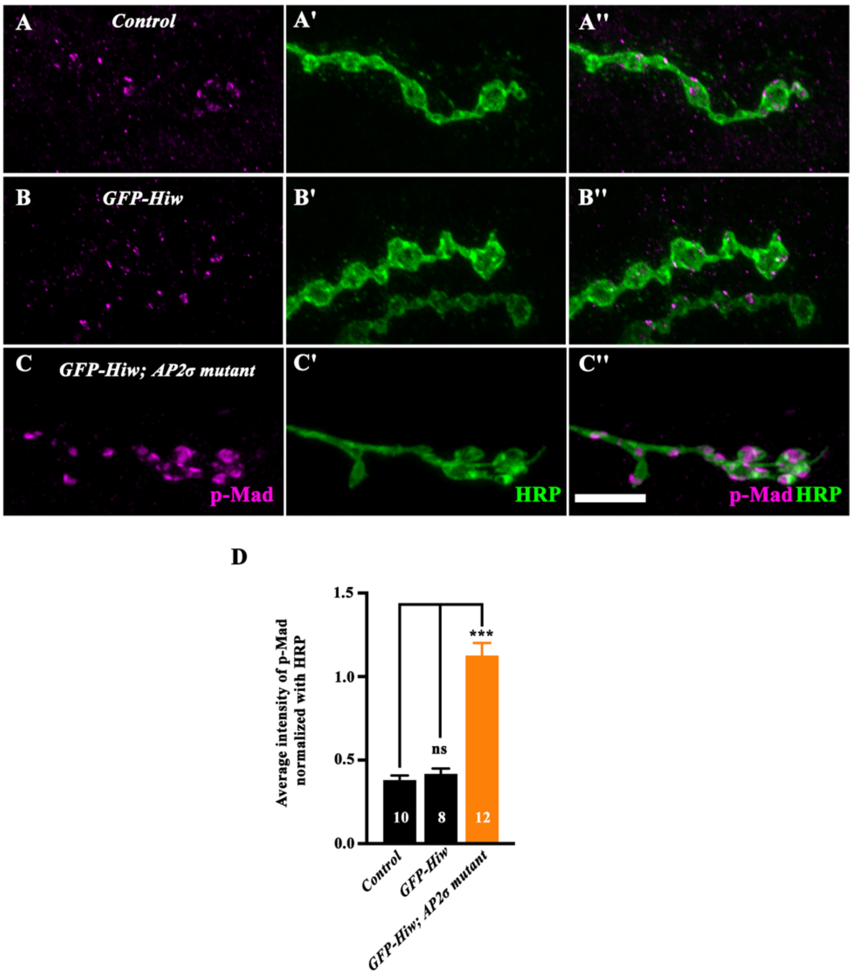


Figure 6 - figure supplement 2

**Figure 6 – figure supplement 2: Endocytic mutants show elevated BMP signaling**

(**A-C''**) Confocal images of muscle 6/7 NMJ co-labeled for pMad (magenta) and HRP (green) in the control (A-A''), *Elav-Gal4> UAS-GFP-Hiw* (B-B''), and *Elav-Gal4>UAS-GFP-Hiw; AP2σ$^{ang7}$/AP2σ-$^{KG}$* (C-C''). The scale bar in C'' represents 10 µm for A-C''. (**D**) Histograms showing the average intensity of p-Mad normalized with HRP at muscle 6/7 NMJ of A2 hemisegment for control (0.38 ± 0.02), *Elav-Gal4> UAS-GFP-Hiw* (0.41 ± 0.03), and *Elav-Gal4>UAS-GFP-Hiw; AP2σ$^{ang7}$/AP2σ-$^{KG}$* (1.13 ± 0.07). At least 8 NMJs of each genotype were used for quantification. The error bars in D represent the standard error of the mean (SEM); the statistical analysis was done using one-way ANOVA followed by post hoc Tukey's test. ***p<0.001, ns: not significant.

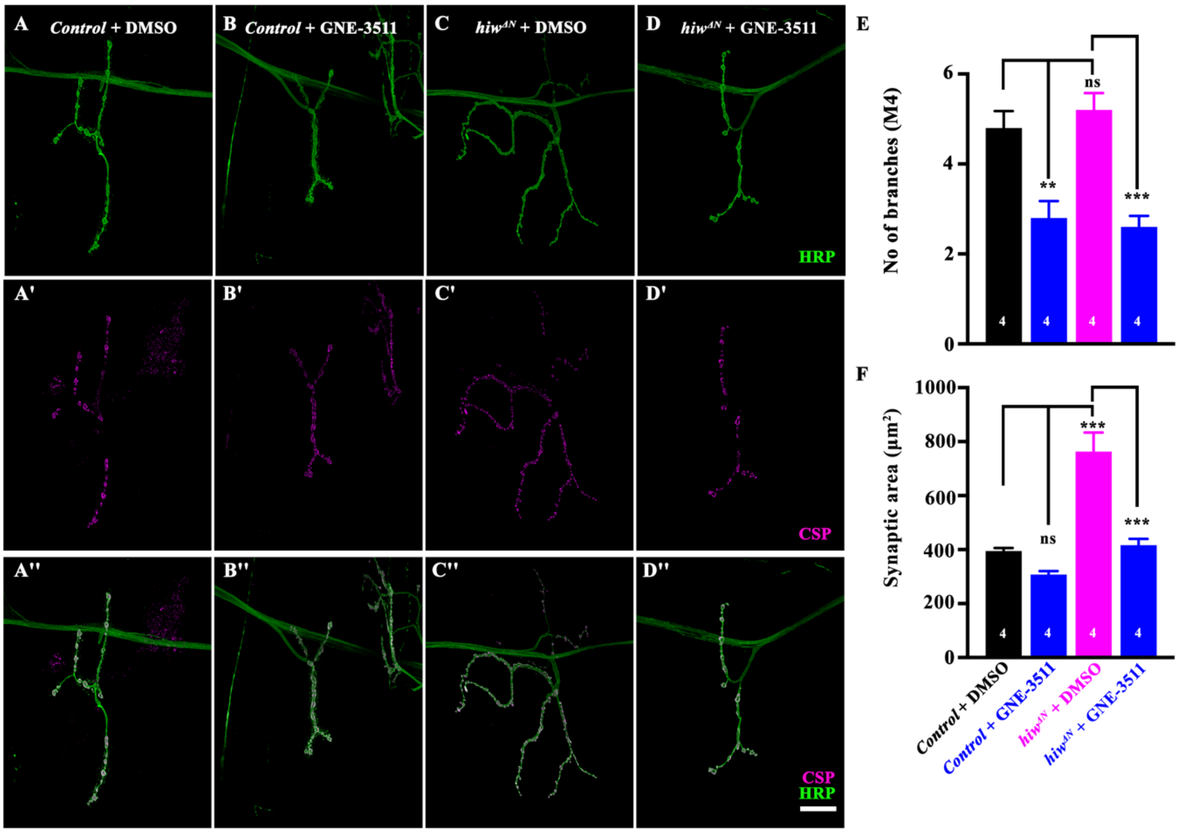


Figure 6 - figure supplement 3

**Figure 6 – figure supplement 3: Treatment with DLKi rescues NMJ overgrowth in highwire mutants**

(**A-D''**) Confocal images of muscle 4 NMJ co-labeled for CSP (magenta) and HRP (green) in the control ($w^{1118}$ + DMSO) (A-A''), $w^{1118}$ + DLKi (B-B''), *hiw*$^{\Delta N}$ + DMSO (C-C''), and *hiw*$^{\Delta N}$ + DLKi (D-D''). The scale bar in D'' represents 10 µm for A-D''.

(**E**) Histogram showing the number of branches at muscle 4 NMJ of A2 hemisegment for control ($w^{1118}$ + DMSO) (4.8 ± 0.37), $w^{1118}$ + DLKi (2.8 ± 0.37), *hiw*$^{\Delta N}$ + DMSO (5.2 ± 0.37), *hiw*$^{\Delta N}$ + DLKi (2.6 ± 0.24). At least 4 NMJs were used for quantification.

(**F**) Histogram showing the synaptic area (µm$^2$) at muscle 4 NMJ of A2 hemisegment for control ($w^{1118}$ + DMSO) (394 ± 11.2), $w^{1118}$ + DLKi (308 ± 12.7), *hiw*$^{\Delta N}$ + DMSO (763 ± 73.8), *hiw*$^{\Delta N}$ + DLKi (416 ± 23.7). At least 4 NMJs were used for quantification. The error bars in E and F represent the standard error of the mean (SEM); the statistical analysis was done using one-way ANOVA followed by post hoc Tukey's test. ***$p<0.001$, **$p<0.01$, ns: not significant.

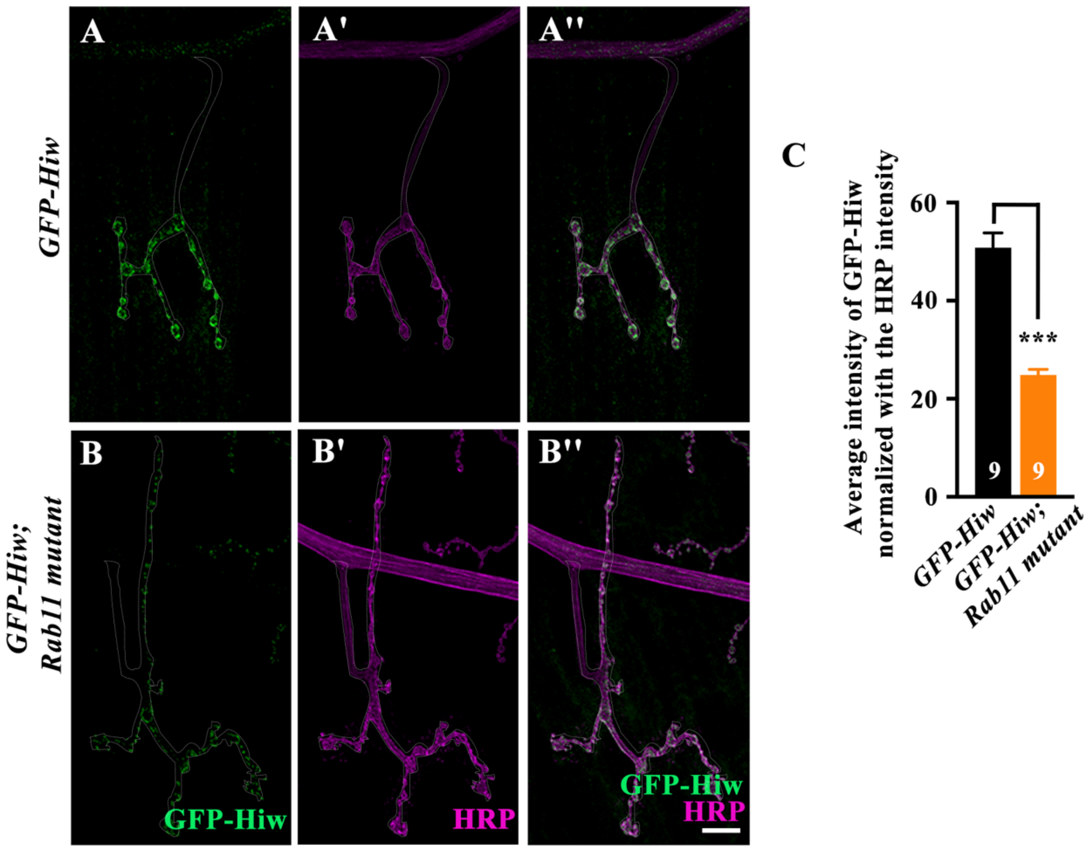


**Figure 7 - figure supplement 1**

**Figure 7 – figure supplement 1: GFP-Hiw levels are reduced at the NMJ in *Rab11* mutants.**

(**A-B''**) Confocal images of muscle 4 NMJ co-labeled for GFP-Hiw (green) and HRP (magenta) in the control (*Elav-Gal4>UAS-GFP-Hiw*) (A-A''), *Elav-Gal4>UAS-GFP-Hiw; Rab11*$^{ex2/93Bi}$ (B-B''). The scale bar in B'' represents 10 µm for A-B''.

(**C**) Histogram showing the average GFP-Hiw levels normalized with HRP in the control (*Elav-Gal4>UAS-GFP-Hiw*) (50.81 ± 3.01) and *Elav-Gal4>UAS-GFP-Hiw; Rab11*$^{ex2/93Bi}$ (24.8 ± 1.13). The error bar represents the standard error of the mean (SEM), and statistical analysis was done using Student's t-test. ***$p < 0.001$. At least 9 NMJs were used for quantification.
